# Functional Mapping of Movement and Speech Using Task-Based Electrophysiological Changes in Stereoelectroencephalography

**DOI:** 10.1101/2024.02.29.582865

**Authors:** Michael A Jensen, Anthony Fine, Panagiotis Kerezoudis, Lily Wong Kisiel, Eva Alden, Dora Hermes, Kai J Miller

**Affiliations:** Department of Neurosurgery, Mayo Clinic, MN, USA; Department of Neurology, Mayo Clinic, MN, USA; Department of Pediatrics, Mayo Clinic, MN, USA; Department of Neuropsychology, Mayo Clinic, MN, USA; Department of Biomedical Engineering, Mayo Clinic, MN, USA

## Abstract

**Introduction:** Stereoelectroencephalography (sEEG) has become the predominant method for intracranial seizure localization. When imaging, semiology, and scalp EEG are not in full agreement or definitively localizing, implanted sEEG recordings are used to test candidate seizure onset zones (SOZs). Discovered SOZs may then be targeted for resection, laser ablation, or neurostimulation. If a SOZ is eloquent, resection and ablation are both contraindicated, so identifying functional representation is crucial for therapeutic decision making.

**Objective:** We present a novel functional brain mapping technique that utilizes task-based electrophysiological changes in sEEG during behavioral tasks and test this in pediatric and adult patients.

**Methods:** sEEG was recorded in twenty patients with epilepsy, aged 6-39 (12 female, 18 of 20 patients < 21 years old), who underwent implanted monitoring to identify seizure onset. Each performed 1) visually cued simple repetitive movements of the hand, foot, or tongue while electromyography was recorded, and 2) simple picture naming or verb generation speech tasks while audio was recorded. Broadband changes in the power spectrum of the sEEG were compared between behavior and rest.

**Results:** Electrophysiological functional mapping of movement and/or speech areas was completed in all 20 patients. Eloquent representation was identified in both cortex and white matter, and generally corresponded to classically described functional anatomic organization as well as other clinical mapping results. Robust maps of brain activity were identified in healthy brain, regions of developmental or acquired structural abnormality, and SOZs.

**Conclusion:** Task based electrophysiological mapping using broadband changes in the sEEG signal reliably identifies movement and speech representation in pediatric and adult epilepsy patients.

## INTRODUCTION

Stereoelectroencephalography (sEEG) has become the most widely used clinical tool for intracranial localization of seizure onset and spread in the treatment of epilepsy^1,2^. It allows physicians to directly characterize the electrical behavior of candidate regions throughout the brain volume during both ictal and interictal periods^3^. During monitoring, electrical stimulation through sEEG may be used to probe sensorimotor, speech & language, vision, or memory function^4,5^. In cases where resection or ablation of epileptogenic tissue is indicated, the spatial overlap of the seizure network with maps of brain eloquence help to determine therapeutic margins such that one can maximize seizure reduction while preserving brain function ^2,6^, especially when dysplastic cortex may carry eloquence^7^. As direct destructive therapy is now being performed through the sEEG leads themselves with radiofrequency ablation^8,9^, extraoperative functional mapping during the monitoring period is essential. Unfortunately, extraoperative stimulation mapping may be limited or impossible when stimulation leads to a seizure or afterdischarges that hinder continuation, particularly in the pediatric brain where the physical properties necessitate higher current^10,11^. Likewise, some patients, especially in the pediatric population^12^, cannot tolerate the MRI scanner environment, preventing pre-implant fMRI.

We describe a technique to complement existing mapping approaches by passively recording electrical signals from sEEG while patients perform simple behavioral tasks at bedside. This task-based electrophysiological (TBE) mapping extends an approach originally developed for brain surface electrocorticography measurements^13-15^, that we initially used in sEEG for the scientific exploration of the brain circuitry of movement^16^. Here we illustrate how TBE mapping using sEEG can be applied in a wide variety of patients and pathologies for the functional mapping of speech and sensorimotor networks.

## MATERIALS AND METHODS

### Ethics statement

The study was conducted according to the guidelines of the Declaration of Helsinki and approved by the Institutional Review Board of the Mayo Clinic (IRB 15-006530), which also authorizes sharing of de-identified data. Each patient or parental guardian provided informed consent as approved by the IRB. All T1 MRI sequences were de-faced prior to publication using an established technique ^17^, to avoid potential identification.

### Subjects

Twenty patients, ages 6-39 (12 female, 18 of 20 patients < 21 years old), participated in our study, each of whom underwent placement of 10-17 sEEG electrode leads for seizure network characterization in the treatment of drug resistant epilepsy (Table 1). Electrode locations were planned by the clinical epilepsy team based on typical semiology, scalp EEG studies, and brain imaging. No plans were modified to accommodate research, nor were extra electrodes added. Nineteen patients participated in our sensorimotor task and eleven participated in the speech task. All experiments were performed in the epilepsy monitoring unit or pediatric intensive care unit at the Mayo Clinic in Rochester, MN.

**Table 1.** Subject Information: Age, Sex, Number of Leads/Laterality, tasks performed (H = hand, T = tongue, F = foot, N = noun, V = verb), seizure onset zones and foci location, hypothesis prior to sEEG, handedness, and speech dominance (if tested), and whether task-based mapping overlapping with a seizure onset zone. Ø indicates that SOZ, which was positive with stim mapping, was lesional

| ID | Age | Sex | # Electrodes/<br># Leads/Side | Tasks | Lesions/ Seizure Onset Zones(s) | Prior Hypothesis | Dominance<br>Hand Speech |  | Task + at<br>SOZ |
| --- | --- | --- | --- | --- | --- | --- | --- | --- | --- |
| S1 | 36 | F | 199/14/B | HTF | **Lesion:** Previous R frontal AVM<br>**SOZ:** No typical seizures recorded | L temporal onset | R | L | N |
| S2 | 16 | F | 217/14/R | HTF | **Lesion:** No lesions<br>**SOZ:** R precentral/postcentral gyri and paracentral lobule | R frontoparietal | L | L | Y |
| S3 | 16 | M | 220/15/B | HTF | **Lesion:** FCD in frontal lobe and postcentral sulcus<br>**SOZ:** Frontal lobe near FCD | R superior frontal/parietal possible FCD | L | L | Y<sup>o</sup> |
| S4 | 16 | F | 223/14/L | HTFNV | **Lesion:** No lesions<br>**SOZs:** Operculum, insula and pre-post central gyri | L insular opercular, left limbic | R | L | Y |
| S5 | 12 | F | 233/17/B | HTFNV | **Lesion:** L parietal lesion and L MTS<br>**SOZ:** L hippocampus and insula | L temporal & bifrontal | L | R | N |
| S6 | 11 | M | 179/12/L | HTFNV | **Lesion:** No lesions<br>**SOZ:** Paracentral lobule | L limbic network, L frontocentral | R | L | Y |
| S7 | 14 | F | 228/15/L | HTFNV | **Lesion:** L inferior frontal at base of gyrus rectus/orbital gyrus cortical thickening and blurring of grey-white junction<br>**SOZ:** L anterior insula/inferior frontal | L temporal, or frontal | R | L | N |
| S8 | 8 | F | 230/17/B | HTFNV | **Lesion:** L parietal FCD<br>**SOZ:** L parietal cortex | L parietal | R | R | N |
| S9 | 39 | M | 185/13/L | HTF | **Lesion:** L MCA Stroke distribution infarct<br>**SOZ:** L hippocampus & precentral gyrus | L central, posterior temporal & parietal | R | L | Y<sup>o</sup> |
| S10 | 15 | M | 185/12/L | HTFNV | **Lesion:** Defect in the upper L cerebral hemisphere contiguous with the L lateral ventricle<br>**SOZ:** Superior-posterior portion of cyst | L perilesional or post central | L | L | Y<sup>o</sup> |
| S11 | 18 | M | 231/14/R | HTFV | **Lesion:** R mesial temporal lobe sclerosis, anterior temporal lobectomy<br>**SOZ:** R posterior temporal lobe and insula | R lesional, perilesional vs. broader temporal limbic | L | R | Y |
| S12 | 15 | F | 159/10/L | HTF | **Lesion:** L mesial temporal lobe sclerosis<br>**SOZ:** L mesial temporal lobe | L temporal, L insular | R | L | N |
| S13 | 13 | M | 196/13/R | HTF | **Lesion:** No lesions<br>**SOZ:** R temporal | R (non-dominant) temporal | R | L | Y |
| S14 | 20 | M | 154/12/R | HTF | **Lesion:** Previous resection in R posterior temporal lobe<br>**SOZs:** 1) Posterior-par-occ-temp junction<br>2) Anterolateral temporal lobe<br>3) Mesial temporal lobe | R temporal lesional/perilesional or broader temporal limbic | R | L | N |
| S15 | 18 | F | 238/15/B | HTF | **Lesion:** L occipital and posteromedial temporal sequelae of Sturge-Weber syndrome.<br>**SOZs:** (15 seizures) L temporo-parieto-occipital region (9), diffuse (5), posterior temporal (1) | L posterior temporal | R | L | N |
| S16 | 7 | F | 201/13/L | NV | **Lesion:** L peri-Rolandic FCD, L frontal gray matter nodular heterotopia<br>**SOZ:** Postcentral sulcus, central sulcus | L parasagittal peri-rolandic | N/A | L | N |
| S17 | 18 | M | 256/16/B | HTFNV | **Lesion:** No Lesions<br>**SOZ:** R mesial temporal | R frontotemporal | R | L | N |
| S18 | 17 | F | 256/17/B | HTFNV | **Lesion:** Bilateral encephalocoele<br>**SOZ:** R temporal head | L temporal | R | L | N |
| S19 | 6 | F | 234/15/L | HTFNV | **Lesion:** Pars intermedia hypointensity, likely Rathke's clef cyst<br>**SOZ:** L mesial temporal lobe | L mesiotemporal and parieto-occipital | R | L | N |
| S20 | 17 | F | 168/13/R | HTF | **Lesion:** Previous R temporal glioma resection<br>**SOZ:** R MTG, insula, hippocampus, amygdala | R temporal | R | L | Y |

### Sensorimotor Task

In the sensorimotor task, three movement types were performed: 1) opening and closing of the hand, 2) side-to-side movement of the tongue with mouth closed, and 3) alternating dorsi- and plantar flexion of the foot (contralateral to the hemisphere of the sEEG montage). Subjects were visually cued to perform these simple self-paced (1 Hz) movements in response to images of a hand, tongue, or foot, and to remain still during interleaved rest periods (blank screen). Twenty cues (trials) of each movement type were shuffled in random order and move/rest trial cues were each presented for 3 seconds (Fig 2). This task was chosen based upon prior work, which has produced clear results in sEEG recordings^16^. The BCI2000 software was used for stimulus presentation and data synchronization^21^. Stimuli were presented on a 53 × 33 cm screen, 80-100 cm from the face. In a minority of trials, patients moved a body part which was not cued (e.g. dorsiflexion when presented with a hand image), but quickly corrected by transitioning to the correct movement. If subject performance was suboptimal, the experimental run would be stopped and re-run later.

**Figure 1.**
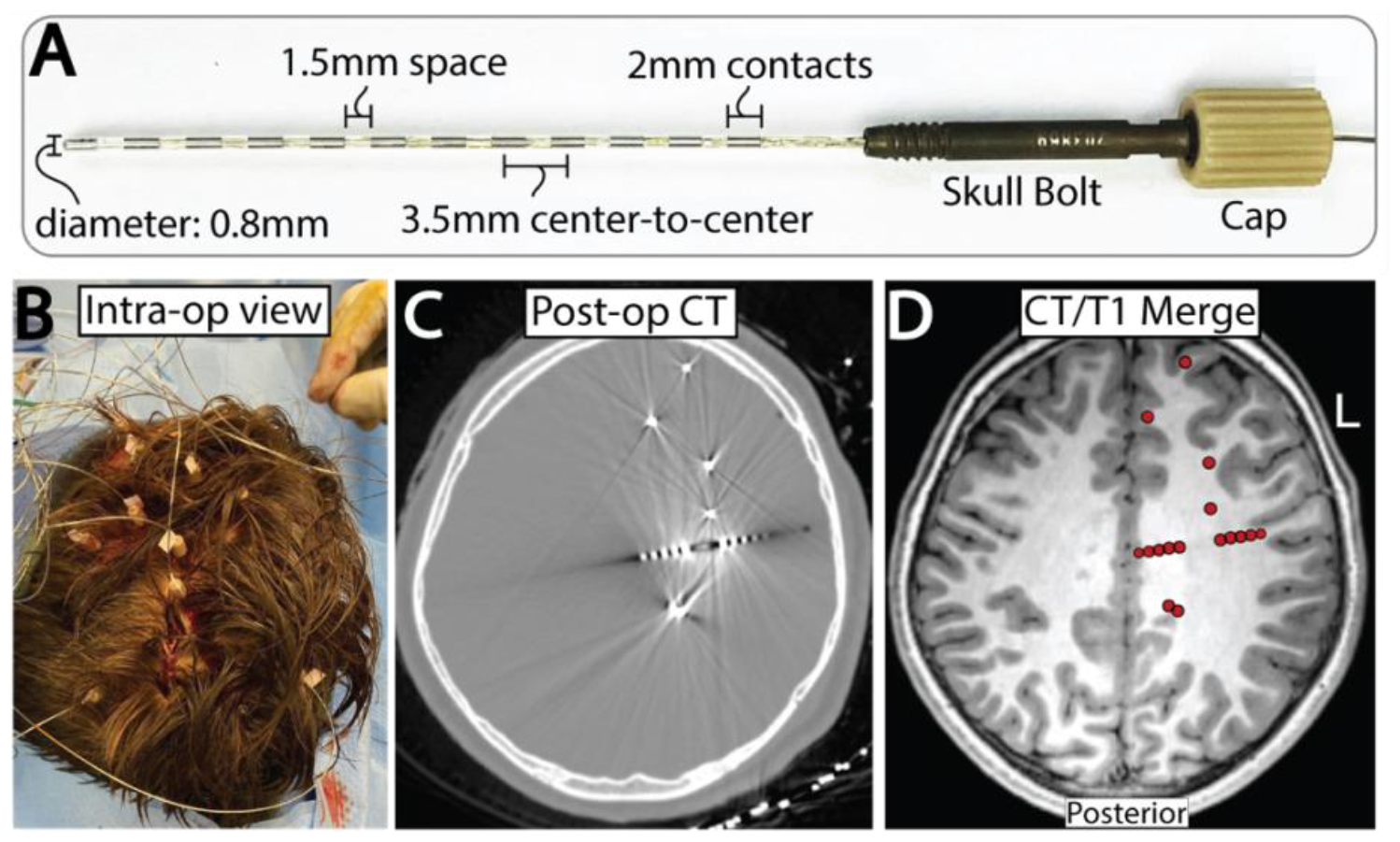
Stereoelectroencephalography (sEEG). **A**. Labeled specifications of an sEEG leads used in clinical practice. **B**. Intraoperative visualization of capped sEEG bolts. **C**. Post operative CT axial slices showing artifact from sEEG contacts. **D**. Merged T1 MRI and CT axial imaging showing electrode locations in the context of its surrounding anatomy.

**Figure 2.**
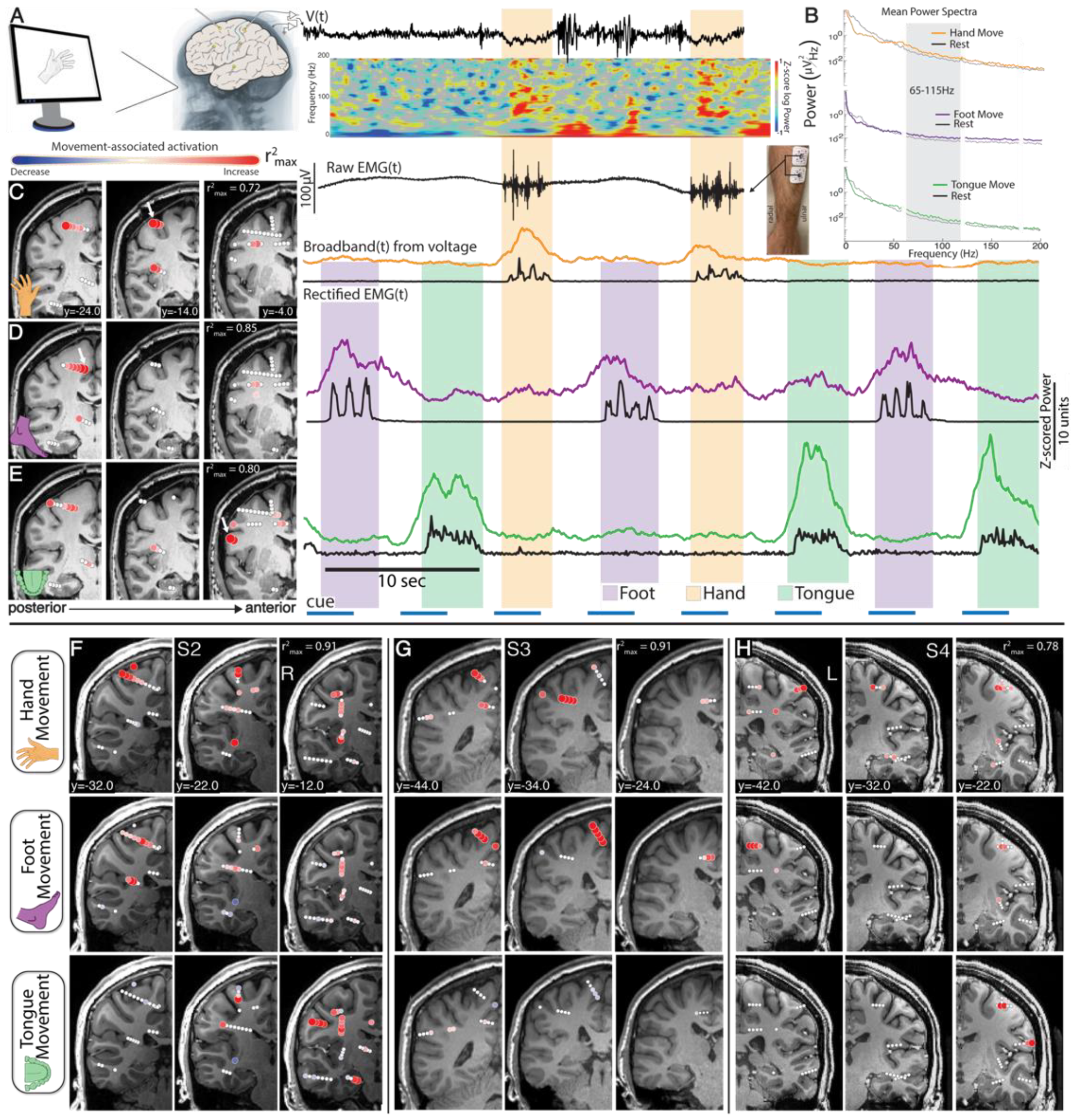
TBE sensorimotor mapping in sEEG – subjects 1-4. **A**. Signals are recorded from all sEEG contacts while the patient moves in response to visual cues displayed on a screen. Bipolar re-referenced channels involved in movement demonstrate broadband increases in power (>50 Hz) during movement periods with a rebound increase in power immediately following movement (<30 Hz) as shown in the time vs. frequency spectrogram. **B**. The mean power spectra of the hand, foot, and tongue channels featured in B-D show movement-related decreases in power within the low frequency bands and increases in broadband power (>50Hz). These plots summarize the spectral dynamics shown in the spectrogram in A but averaged across all rest and movement periods. **C-E**. *r*^*2*^ activity maps: subjects perform contralateral movement tasks consisting of opening and closing their hand (B), dorsi and plantarflexing their foot (C), and moving their tongue laterally with mouth closed (D) in response to visual cues while recording surface EMG. Somatotopically selective channels are revealed by high frequency broadband power (65-115 Hz) increases during movement. White arrows in the T1 coronal slices (10 mm thick) indicate channels corresponding to the broadband power time series shown (right). F-H. As in C-E.

Electromyography (EMG) was measured from the forearm flexors/extensors (hand), base of chin (tongue), and anterior tibialis (foot) during the sensorimotor task (Fig 2). All sEEG and electromyographic (EMG) signals were measured in parallel, and delivered to the research DC amplifier (g.HIAmp system, gTec). sEEG and EMG signals were synchronized with visual stimuli using the BCI2000 software^21^. Data were segmented into movement and rest periods using synchronized EMG as described previously^16^.

### Speech Tasks

In both of our two speech tasks, subjects were presented with a visual cue for 3.5 sec (e.g. Picture of a soccer ball), and they either stated a self generated noun (e.g. “ball”) in the ‘noun production task’ or a self generated verb associated with the picture (e.g. “kick”) in the ‘verb generation task’. Both the noun production and verb generation tasks used the same stimuli (Fig 3, 36 stimuli). Subjects were instructed to remain silent when no stimulus was present (2 sec rest). The BCI2000 software was used for stimulus presentation and data synchronization^21^, with stimuli presented on a 53 × 33 cm screen, at a distance 80-100 cm from the face. If subjects were not participating with the task, the experimental run would be stopped and re-run later. Data were segmented into speech and silent periods based on synchronized audio recorded during the task. To account for the neural activity in the brain preceding phonation, 500 ms of data prior to audio-detected speech were included in each speech epoch. Although these tasks function to identify speech representation, engagement of language network regions not directly involved in speech motor output is possible. Conceptually, speech refers to a motor function and language is a cognitive process, thus completing these tasks has the potential to elicit activity in regions supporting both functions.

**Figure 3.**
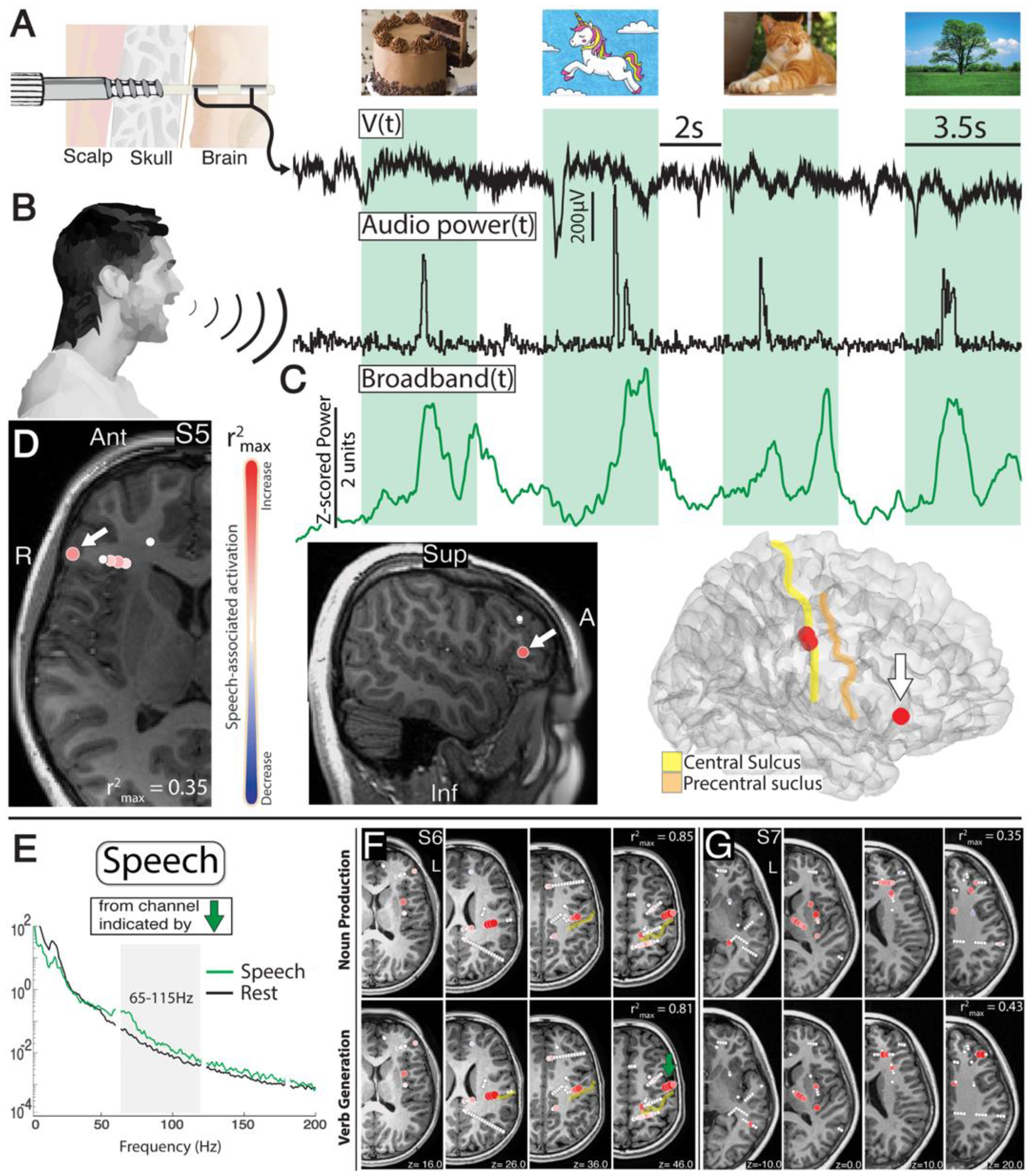
TBE speech mapping in sEEG– subjects 5-7. **A**. sEEG recorded during speech tasks in a bipolar fashion. **B**. Subjects were shown images for 3.5 sec (green background). They either named the images (noun, e.g. “cake”), or stated and action associated with the image (verb, .e.g “eat”). **C**. Broadband time series from channels recording in Broca’s area (white arrow in D) show modulation of local cortical activity during speech (verb generation). **D**. *r*^*2*^ activity maps shown in an axial slice, sagittal slice, and brain rendering. Note: Subject 5 was right language dominant (Fig 9). **E**. Speech vs rest mean power spectra during the verb generation task shows speech-related decreases in power within the low frequency bands and increases in broadband power (>50Hz) taken from the channel in E (green arrow). **F-G**. Axial (10 mm thick) *r*^*2*^ activity maps generated during verb generation and noun production tasks in subjects 6 and 7.

### Stereoelectroencephalography Recordings

Intracranial sEEG signals were recorded relative to a clinician-selected reference in the white matter a-priori believed to be far away from tissue likely to have seizure involvement. Voltage timeseries were recorded with the 256-channel g.HiAmp amplifier (gTec, Graz, Austria). Recordings were sampled at 1200Hz, with an anti-aliasing filter, which dampened the signal by 3dB at 225Hz.

### Lead Placement, Electrode Localization, and Re-referencing

The platinum depth electrode contacts (DIXI Medical) were 0.8mm in diameter with 2mm length circumferential contacts separated by 1.5mm (Fig 1). Each lead contained 10-18 electrode contacts. Surgical targeting and implantation were performed in the standard clinical fashion^2^. Intraoperatively, anchoring bolts were placed into stereotactically aligned 2.3 mm holes in the skull and then leads were advanced to target through the bolts (Fig 1).

Electrode-to-anatomic localizations were determined by co-registration of post-implant CT scan to pre-implant MRI (Fig 1). Each preoperative T1 MRI was realigned to the anterior and posterior commissure stereotactic space (AC-PC)^18^, and then co-registered to the post-implant CT using SPM12^19^.

All data were re-referenced in a bipolar fashion, producing channels that reflect mixed activity between voltage time series measured at two adjacent electrode recording sites (Figs 2a, 3a). Plotted points for brain activity in this study each represent an interpolated point between the two electrodes that make up each differential pair channel. Only adjacent differential pair channels were considered - i.e. 1.5mm from one another, on the same lead. In each activity map, channels were plotted using the SEEGVIEW package, which slices brain renderings, and projects channels to the closest slice^20^. The purpose of this tool is to present analyses in an interpretable, clinically familiar manner. Projection distance increases alongside slice thickness, and a channel reflecting activity in the gray matter may artifactually appear to be in white matter after projection to the nearest slice. Thus, there is a tradeoff between the number of channels that can be visualized and the accuracy of representation for each channel’s anatomical position.

### Signal Processing and Analysis

Trial-by-trial power spectral analysis were performed using custom scripts in MATLAB. For each movement or speech trial, averaged power spectral densities (PSDs) were calculated from 1 to 300 Hz every 1 Hz using Welch’s averaged periodogram method, with 1 second Hann windows to attenuate edge effects and 0.5 second overlap^22^. The averaged PSD for each movement/speech or rest trial was normalized to the global mean across the entire experimental run (note that movement and speech tasks were all performed on different experimental runs). The PSDs were normalized in this way because brain signals of this type generally follow a 1/f, power law shape^23^ - lower frequency features dominate in the absence of normalization. From each of these normalized single trial PSDs, averaged power in a broadband high frequency band (65-115 Hz) was calculated for subsequent analysis, as previously described^16^. This captures broadband activity above the known range of most oscillations that might contaminate it, and avoids ambient line noise at 60 Hz and 120 Hz.

For each bipolar re-referenced channel, we calculated separate signed *r*^*2*^ cross-correlation values (r^2^) of the mean spectra from 65-115 Hz for each movement modality. Each channel’s *r*^*2*^ value was determined by comparing mean power spectra between movement/speech trials (separately) and rest. To minimize the cross-effects of movement-specific rebound effects^24^, movement trials of each type were only compared with rest trials that followed that same movement type:

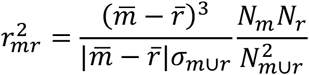

where m denotes power samples from movement/speech, r denotes samples from rest, and 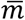 and 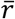 denote sample mean. m_⋃_r represents the union of power samples in movement/speech and rest periods. N_m_ and N_r_ denote the total number of rest and movement/speech samples and N_m∪r_ = N_m_ +N_r_. Thus, *r*^*2*^ is the percentage of the variance in m that can be explained by a difference between the individual means in the subdistributions, 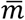 and 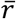 ^25^. The sign indicates whether power is increasing or decreasing with behavior, as illustrated by red and blue circles in each activity map (Figs 2-8, Supp Figs 2-26). We used a two-sample t-test to calculate the significance of the difference between high frequency band power during movement vs. rest epochs. Channels are colored only if the uncorrected behavior versus rest p-value was less than 0.05. All insignificant channels were plotted as a white circle of fixed diameter.

### Clinical Stimulation Mapping

Stimulation was delivered in a bipolar fashion with a 1Hz (in some cases of language mapping) or 50 Hz pulse frequency, 200 ms pulse duration, 2-5 second train duration, and stimulation intensities from 1 to 5 mA (Supp Fig 1). During stimulation, patients were monitored for fasciculations and muscle jerks, abnormal movement interruptions, speech production, or abnormal speech arrest/delay by a team of neurologists and EEG technicians. Any sensations experienced by the patients were communicated in real-time to the team and documented immediately. All observed and patient-reported responses to stimulation were considered positive stimulation results. If no observed or subjective responses occurred up to 5 mA intensity, stimulation progressed to the subsequent electrode pair. Afterdischarges (ADs) were recorded, and based on their duration and location, the neurology team determined whether to continue with further stimulation at the same site. If stimulation induced a clinical seizure, it was terminated. In the case stimulation induced a very short-lasting electrophysiologic seizure, the clinical decision to continue stimulation was based on patient symptoms.

### Interpretation of Stimulation Results

The results of all stimulation mapping sessions are recorded in their electronic medical record (EMR). Authors MAJ and AF categorized stimulation responses. Sensorimotor responses were considered positive when the response descriptions in the EMR contained the following terms: “aphasia”, “activation of [insert muscle]”, “puckering”, “motor pause”, “tingling”, “speech arrest/pause/impairment”, “behavioral arrest”, “wave of sensation”, “shaking”, “twitch”, “[insert appendage] weakness”, “[insert muscle] extension/flexion/pronation/supination. For strict sensorimotor interpretations (Supp Fig 25), either speech arrest responses were removed or only the responses involving muscle activation/fasciculation/twitching/weakness were included. Speech responses were considered positive when the response descriptions in the EMR contained the following terms: “aphasia”, “speech arrest/pause/impairment”, “slowing of reaction when naming”.

### Data availability

All data & code needed to reproduce findings are publicly available on Open Science Framework (DOI 10.17605/OSF.IO/B7N5C).

### Code availability

All code necessary to reproduce findings are pub licly available on GitHub (https://github.com/michaeljensen42/TBE-mapping-in-sEEG).

## RESULTS

sEEG-based TBE mapping allows for volumetric assessment of function by simultaneously recording activity from all intracranial channels (Fig 1). Sensorimotor, noun production, and verb generation tasks were performed in a primarily pediatric cohort down to 6 years of age without significant challenges (Table 1). Volitional movement typically began 0.5 to 1 second after stimulus cue presentation and ended at the beginning the rest period. Concurrent surface EMG recordings were synchronized with sEEG signal to allow for accurate segmentation of data into true movement and rest trials (Fig 2). Speech was often produced in the middle of the stimulus period, and audio recordings synchronized to sEEG signal were used to accurately segment data into speech and rest trials (Fig 3). High frequency broadband power localizes activity to both primary and association sensorimotor and speech cortical regions across patients and is easily interpretable when viewed on brain slices (Fig 2b-h, Fig 3c-f, Supp Figs 2-24). The sensorimotor task commonly evoked activity in the primary sensorimotor, supplementary motor, and pre-motor regions (Figs 2, 4, Supp Figs 2-16), while speech tasks evoked activity in pre, primary, and cingulate motor tongue regions in addition to superior temporal (Wernicke’s area), supramarginal, and inferior frontal (Broca’s area) gyri due to the inevitable engagement of language circuitry (Figure 3, 4, Supp Figs 17-24). Primary cortical regions consistently demonstrated greater behaviorally induced broadband power increases than non-primary regions (Figure 2-4).

**Figure 4.**
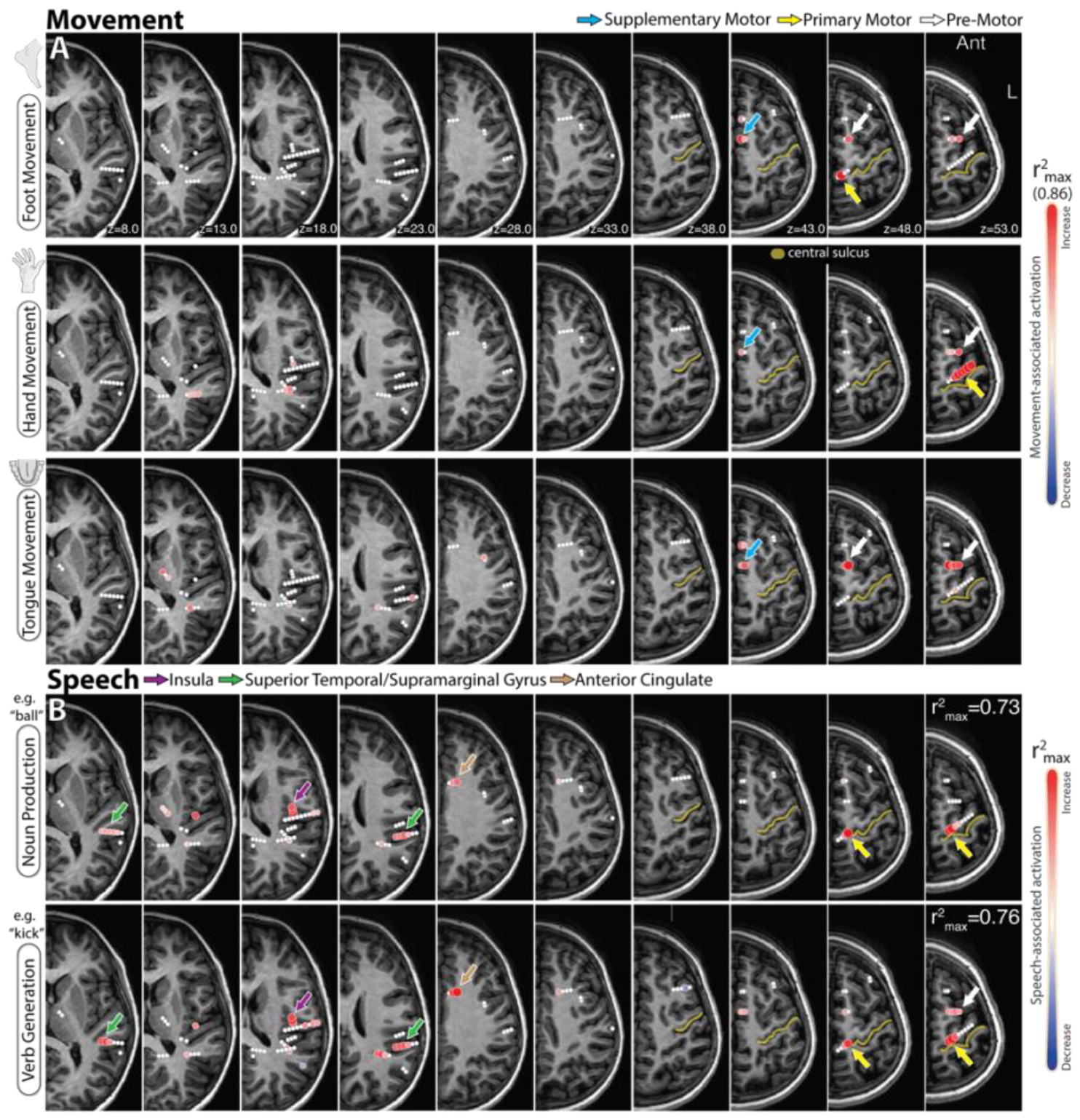
Activity maps across all modalities in subject 8. **A-C**. T1 axial slices (5 mm thick) demonstrating *r*^*2*^ activity maps generated during EMG-defined hand (A), tongue (B), foot (C) movement. r^2^ values are scaled to the maximum across all three movement types. Somatotopically organized activity is observed in expected locations (colored arrows). **B**. As in A, but during noun and verb tasks. As speech tasks were run separately, *r*^*2*^ values are displayed for each task.

TBE mapping is reliable, localizing function within healthy (Fig 2, 3, 4, 8), injured, and congenitally abnormal tissue (Fig 6,7) whether or not the tissue was epileptogenic (Fig 4, 5). For example, TBE effectively localized function in the context of 1) non-lesional seizure onset zones (Fig 5), 2) encephalomalacia in an adult three years after MCA stroke (Figs 6a,b, Supp Fig 5), 3) a porencephalic cyst of an adolescent who suffered a perinatal hemorrhagic stroke (Fig 6c,d, Supp Fig 6), and 4) focal cortical dysplasia (Fig 7). Interestingly, TBE mapping also identified white matter activity during sensorimotor (Supp Fig 25) and speech (Supp Fig 26) tasks, though sparsely, and in behaviorally relevant tracts. This may be simply reflective of true white matter activity and the depolarization from passing action potentials at the nodes of Ranvier, but it may also be due to volume conduction from nearby cortex or field potentials from glial tissue^26^.

**Figure 5.**
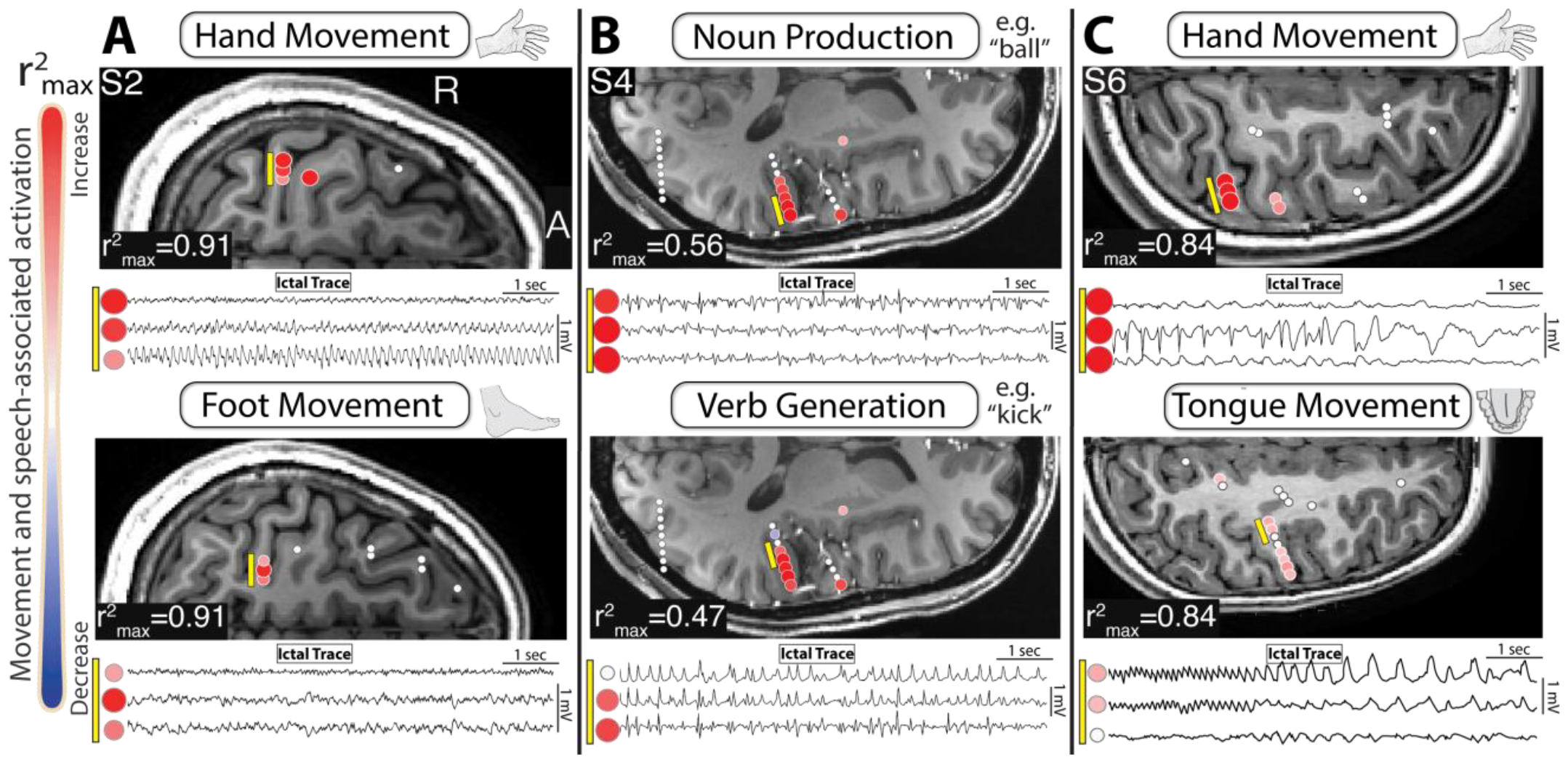
TBE mapping identifies function in non-lesional seizure onset zones – subjects 2, 4, 6. **A**. Robust hand (top) and foot (bottom) activity maps within epileptogenic channels (white arrows). Corresponding ictal traces are shown below for subject 2. **B**. As in A for subject 4 during noun (top) and verb (bottom) speech tasks. **C**. As in A for subject 6 during hand (top) and tongue (bottom) movement. **Note:** Ictal traces were identified by epileptologist (A.F.)

**Figure 6.**
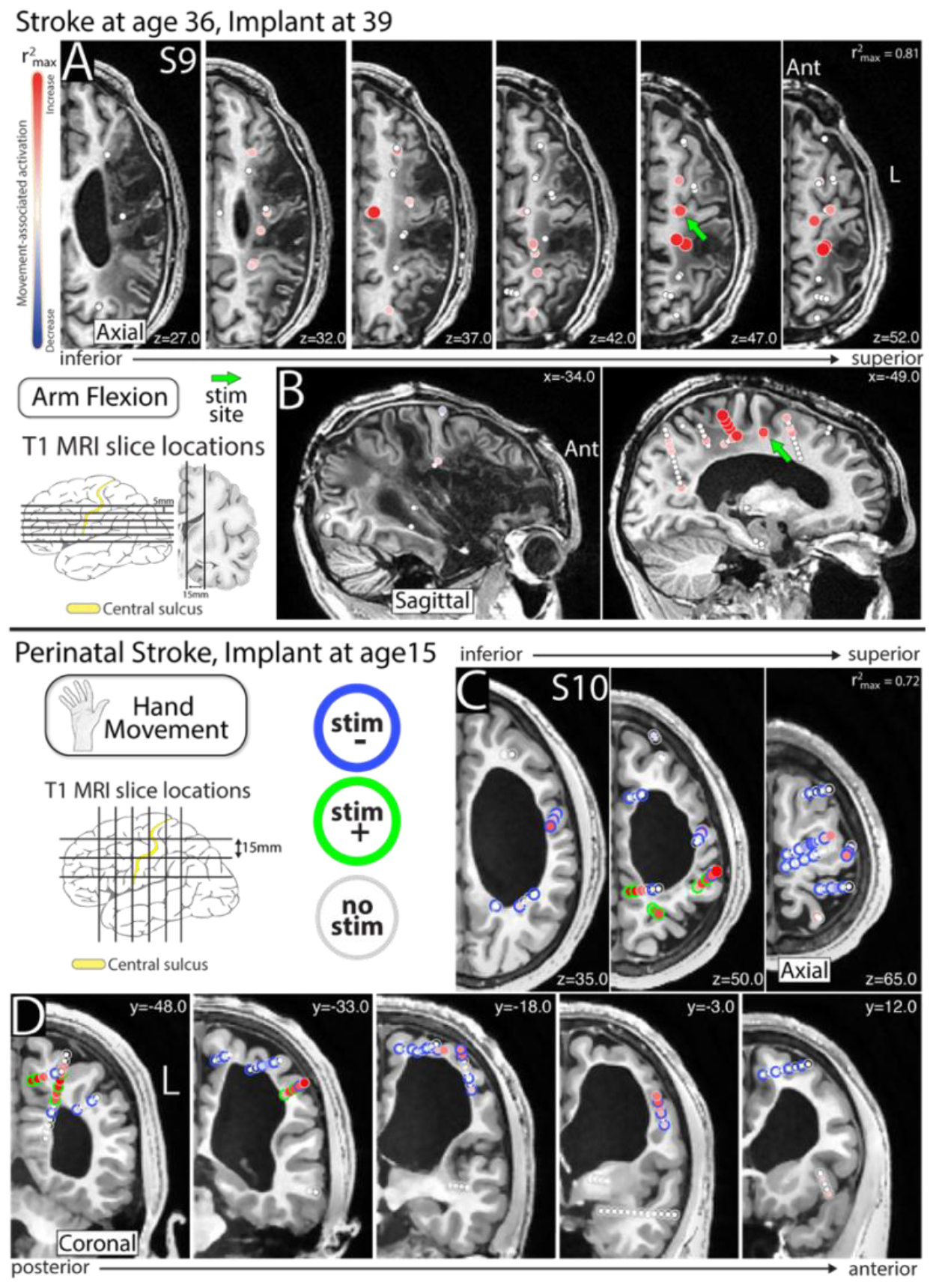
TBE mapping identifies function within injured brain in patients with previous stroke - subjects 9,10. **A**. A 36 year old man (S9) developed seizures after a dominant, left-sided, MCA stroke, and was implanted with sEEG 3 years later. Robust movement (arm flexion) activation was observed in injured tissue at the margin of encephalomalacia (shown in axial 5 mm thick *r*^*2*^ activity maps). Clinical stimulation was performed at a single site (green arrow), leading to a sensorimotor response in the right arm. Five minutes later stimulation was repeated at the same site with higher amplitude, and a generalized seizure ensued. **B**. As in A, but with sagittal slices (15 mm thick). **C**. A 15 year old boy (S10) developed seizures after a perinatal hemorrhagic stroke. Hand movement activation around the site of stroke is shown in axial (15 mm thick) *r*^*2*^ activity maps. Clinical stimulation was performed, and outer rings indicate stimulation results (stimulation negative – blue, stimulation positive – green, no stimulation – gray). Note: For this patient, all positive stimulation sites were post-central rather than motor in nature. **D**. As in A, but with coronal (15 mm thick) slices.

**Figure 7.**
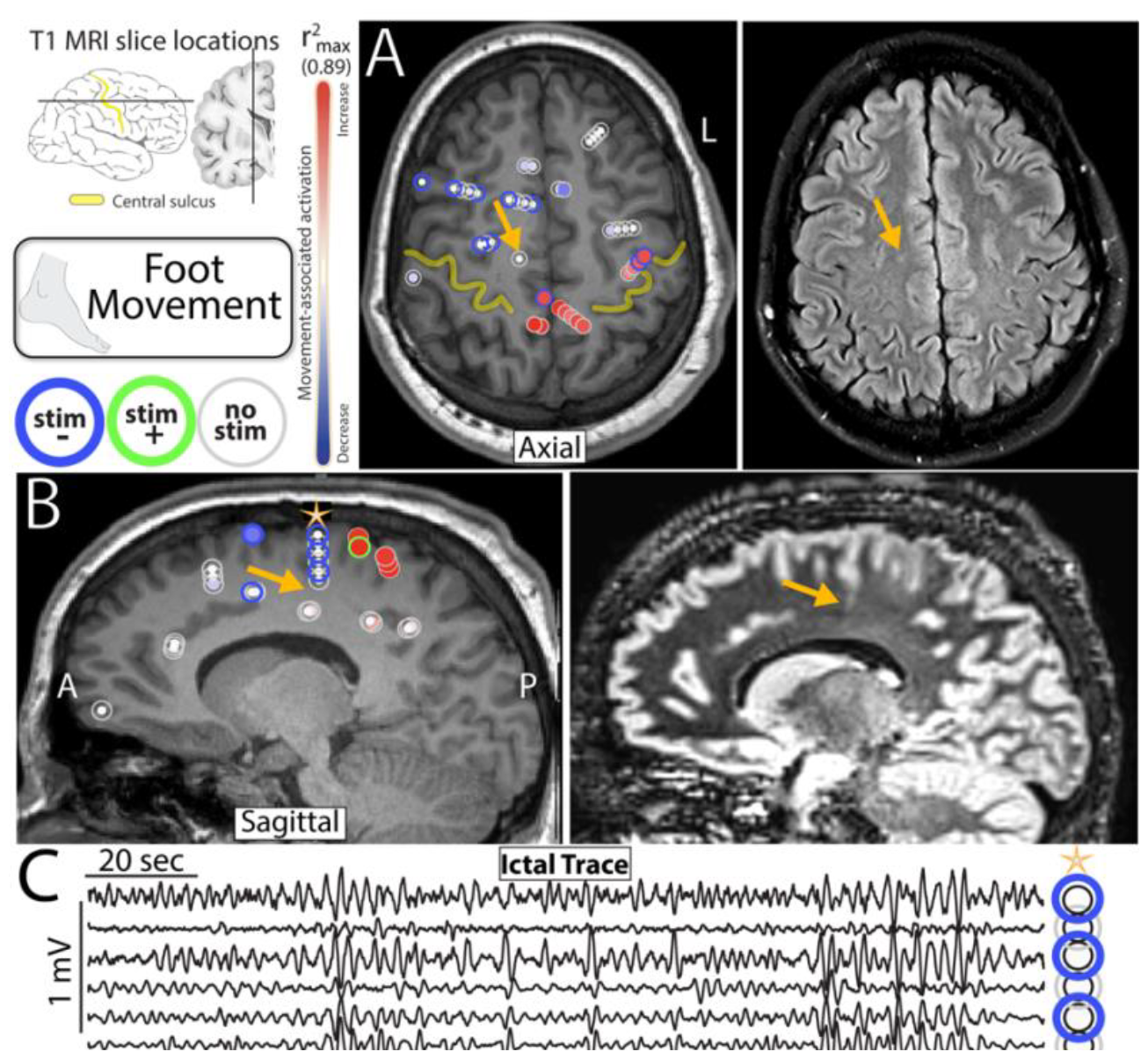
TBE sensorimotor mapping identifies function adjacent, but not within, a focal cortical dysplasia - subject 3. **A**. T1 (left) and T2 FLAIR (right) showing focal cortical dysplasia (FCD – orange arrow). Foot movement representation around the FCD are shown as in Figure 5 C,D. **B**. As in A, for sagittal slice. **C**. Voltage traces taken from the six channels near the FCD highlighted by a star in B during a typical seizure. Note: motor representation was not found by neither stimulation nor TBE mapping in the focal cortical dysplasia. This is in stark contrast to the activity within homologous (premotor/SMA) sites in the right most slices of Figure 4a.

**Figure 8.**
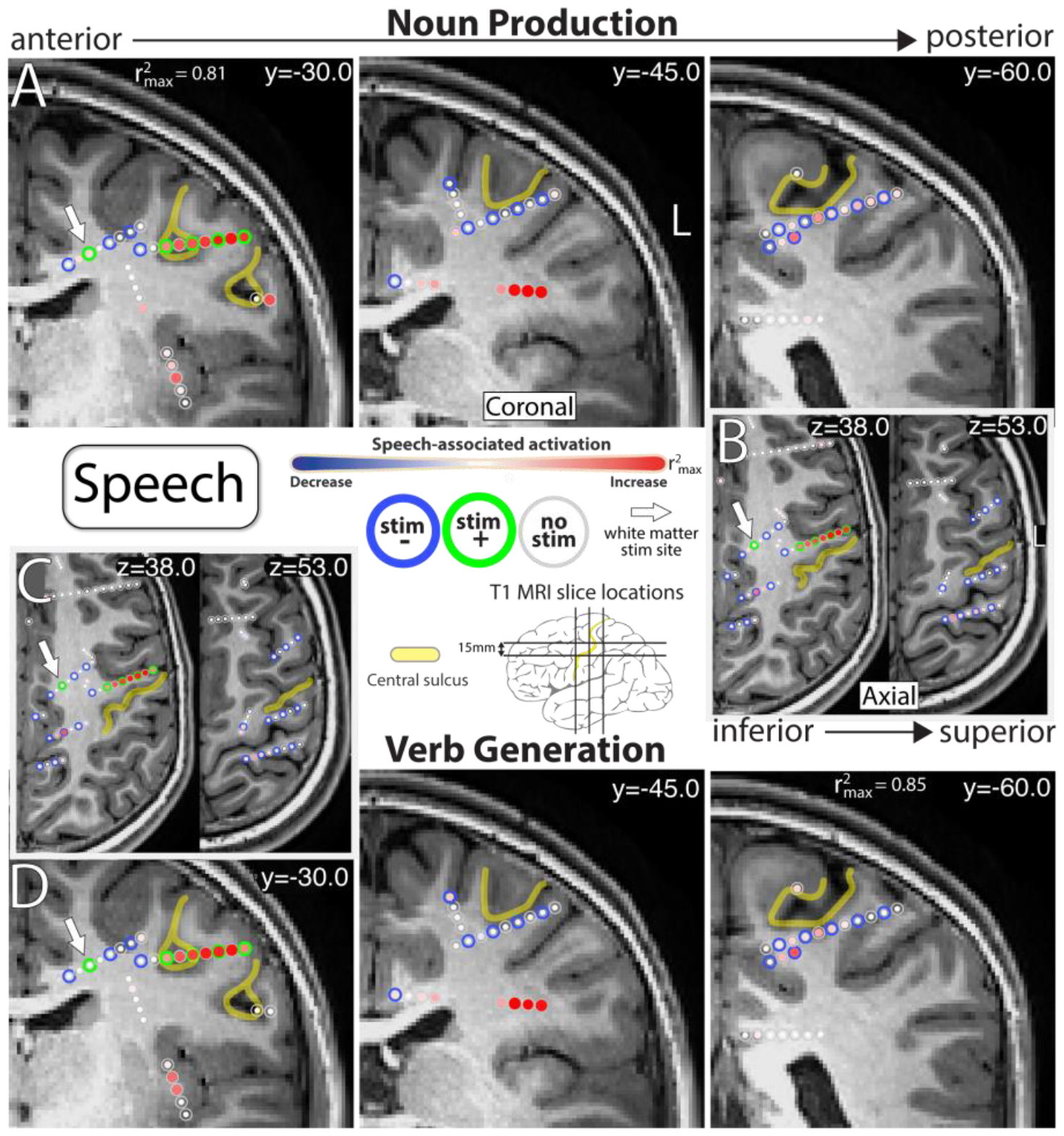
Comparison of TBE and Stimulation Mapping During Speech Tasks - Subject 6. **A**. Activation associated from noun production is shown in coronal (15 mm thick) *r*^*2*^ activity maps. Simulation results are shown as in Figure 5 C,D. **B**. As in A, but with axial slices. **C**. As in B, but during verb generation. **D**. As in A, but during verb generation. Yellow highlighting approximates the location of the central sulcus. Note: white arrow indicates positive stimulation site located in white matter.

There is general agreement between TBE and stimulation mapping. This was robust in the pathological margins of injured brain (Fig 6), focal cortical dysplasia (Fig 7), and within healthy tissue (Fig 8). When disagreement is present, it is typically seen as a site with broadband power increases but no response to stimulation (stimulation -, TBE +, Fig 6c,d). Conversely, of all sites in which stimulation led to positive sensorimotor or speech responses, 99% and 96% were also positively identified (p < 0.05) using sensorimotor and speech TBE mapping respectively (Supp Fig 27, 28). If sensorimotor stimulation results are interpreted more strictly (Supp Fig 25), agreement was 100% (“stricter” interpretations of speech results were not possible without introduction of bias due to intrinsic difficulty). The opposite discordance (stimulation +, TBE -) was seen only rarely, and in the context of white matter stimulation (Fig 8). The fMRI BOLD signal has been shown to be well correlated to broadband power at the brain surface^27^, and we find anecdotally that clinical fMRI speech and movement tasks correspond well to TBE mapping (Fig 9).

**Figure 9.**
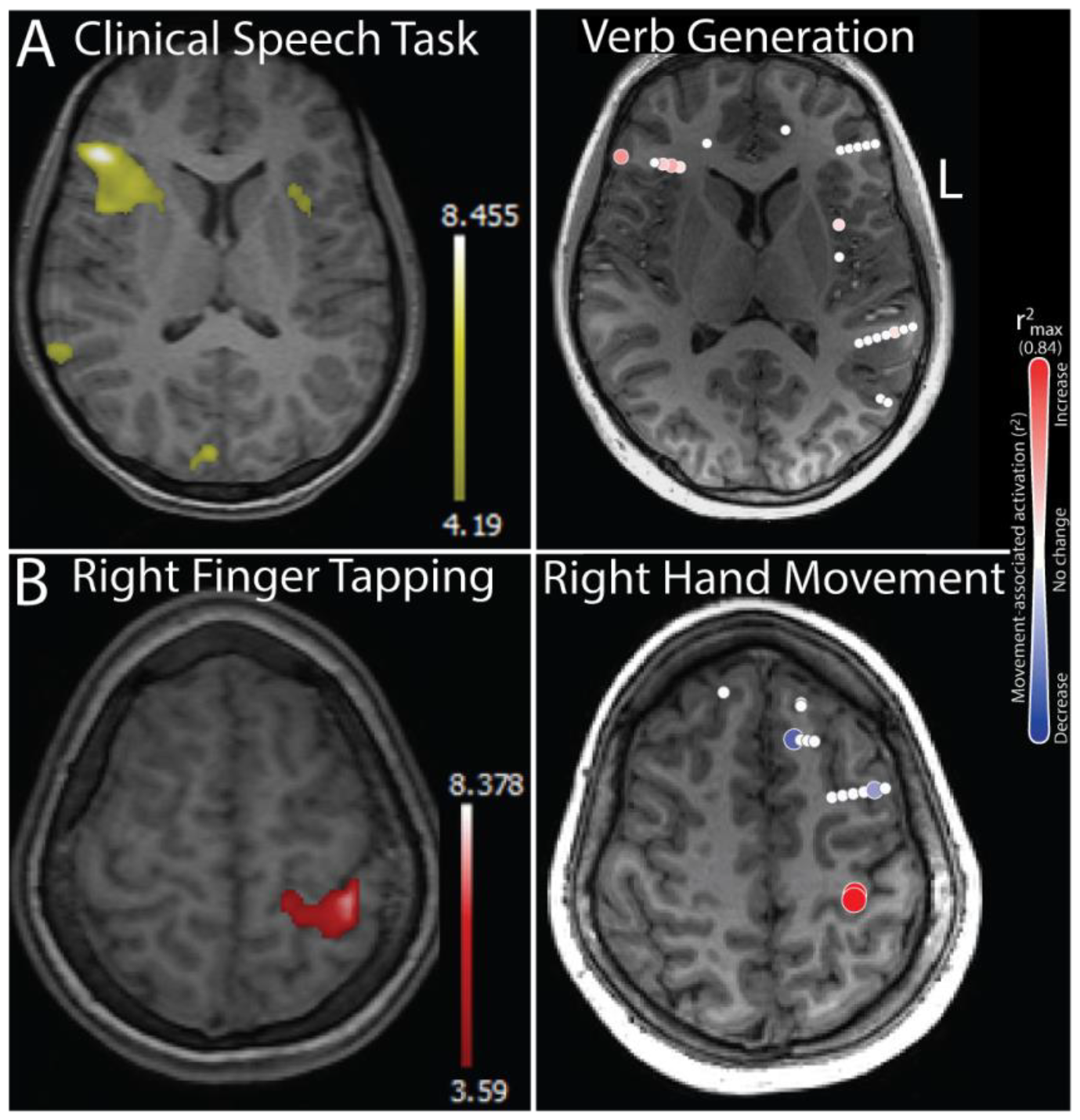
Comparison of Clinical fMRI and sEEG Functional Mapping Results - Subject 5. **A**. sEEG speech mapping agrees with fMRI clinical speech mapping. Note: subject is right language dominant. **B**. sEEG hand movement mapping correspond to clinical fMRI finger tapping task.

## DISCUSSION

Epilepsy impacts a heterogenous population with a diverse set of pathologies, so the clinical mapping tools physicians use should be simple to apply and effective across a wide variety of contexts. We have shown that sensorimotor and speech task based electrophysiological mapping using sEEG can be conducted in patients as young as six years old and allows for robust localization of function within both healthy and pathologic tissue.

Although we do have an existing set of clinical tools for functional brain mapping, each individual patient will often require an overlapping tableau of different functional localization techniques that may include fMRI, electrical stimulation mapping^28^, MEG^29^, TMS^30^, and function inferred by the behavioral semiology induced at the time of seizure onset. Each tool has its own limitations, and it is rare that a single modality provides sufficient confidence for a resectional margin. Electrical stimulation may result in afterdischarges or a seizure (Fig 6a,b), hindering complete mapping^28^, and some patients, especially pediatric, may not be able to tolerate the MRI scanner environment^12^. Moreover, electrical stimulation with sEEG may affect white matter connections^4^, potentially complicating the interpretation of the region and network involved in function. This TBE mapping approach provides an additional tool to complement the existing repertoire. It has the advantage that it may be performed at the bedside in a single session, and that variable compliance by the patient (as may often be the case in pediatric patients) may be adjusted for by careful segmentation of the behavior measured from the EMG or the recorded audio.

As is the case when reconciling fMRI with electrical stimulation mapping, interpreting discrepancies between TBE mapping and stimulation should allow for nuance. Functional networks, such as those governing speech or movement, often contain redundancy across cortical regions which can substitute for one another to sustain behavior despite focal interruption^31^. This redundancy is a common feature of association cortices^31^, and less likely in primary cortex^16^. Such shared representation is the most probable explanation for channels that demonstrate broadband power increases (TBE +) during behavior but no obvious response to stimulation (Supp Fig 29). This helps to explain the discordance seen in Figure 6C&D, in which hand representation has likely been taken over by the hemisphere ipsilateral to movement. In the exceptionally rare case where electrical stimulation of a TBE negative site elicits a clinical response, we have found it to be most likely due to 1) white matter stimulation (Fig 8, white arrow) which can activate distance cortex^28,32^, 2) a lack of pre-stimulation behavioral baselining (Supp Fig 29), or 3) over-interpretation of an ambiguous clinical state (an incorrect behavioral attribution).

What has been demonstrated here for speech and movement may be extended to any functional modality depending upon clinical need. Therapeutic intervention for epilepsy should explicitly preserve memory, cognition, and vision to the greatest degree possible - this TBE mapping approach is immediately generalizable to these contexts.

## CONCLUSION

Task-based electrophysiological mapping using broadband changes in the sEEG signal localizes movement and speech representation accurately and reliably across behaviors, patient populations, and tissue quality. It is a robust tool that can be used at the bedside to complement existing techniques and improve decision making and intervention for patients with epilepsy.

## ACKNOWLEDGEMENTS

We wish to extend our deepest gratitude to the patients who volunteered their time to participate in this research, to Bambi Wessel, Cindy Nelson, Lidya Legesse, the staff at St. Mary’s hospital, and Peter Brunner for his critical assistance in the assembly of our electrophysiology rig and operations of the BCI2000 software. This work was supported by the NIH-NCATS CTSA KL2 TR002379 (KJM), and by the Brain Research Foundation with a Fay/Frank Seed Grant (KJM), NIH P41-EB018783 (PB), NIH U01-NS128612 (KJM, PB, GAW), NIH R01-EB026439 (PB), NIH U24-NS109103 (PB), NIH U01-NS108916 (PB). The contents of this manuscript are solely the responsibility of the authors and do not necessarily represent the official views of the NIH. Our funders did not play a role in study design, data collection and analysis, decision to publish, or preparation of the manuscript.

