## supplementary material for "Functional Mapping of Movement and Speech Using Task-Based Electrophysiological Changes in Stereoelectroencephalography"

**Figure 1:** Schematic Description of Cortical Stimulation Mapping (Page S2)

**Reproduction of Motor Maps Across Subjects Not Shown in Main Article**

**Figure 2:** Task-based Electrophysiology (TBE) Maps During Hand, Tongue, & Foot Movement for Subject 5 (Page S3)

**Figure 3:** TBE Maps During Hand, Tongue, & Foot Movement for Subject 6 (Page S4)

**Figure 4:** TBE Maps During Hand, Tongue, & Foot Movement for Subject 7 (Page S5)

**Figure 5:** TBE Maps During Arm Flexion for Subject 9 (Page S6)

**Figure 6:** TBE Maps During Hand, Tongue, & Foot Movement for Subject 10 (Page S7)

**Figure 7:** TBE Maps During Hand, Tongue, & Foot Movement for Subject 11 (Page S8)

**Figure 8:** TBE Maps During Hand, Tongue, & Foot Movement for Subject 12 (Page S9)

**Figure 9:** TBE Maps During Hand, Tongue, & Foot Movement for Subject 13 (Page S10)

**Figure 10:** TBE Maps During Hand, Tongue, & Foot Movement for Subject 14 (Page S11)

**Figure 11:** TBE Maps During Hand, Tongue, & Foot Movement for Subject 15 (Page S12)

**Figure 12:** TBE Maps During Hand, Tongue, & Foot Movement for Subject 17 (Page S13)

**Figure 13:** TBE Maps During Right Hand, Tongue, & Foot Movement for Subject 18 (Page S14)

**Figure 14:** TBE Maps During Left Hand & Foot Movement for Subject 18 (Page S15)

**Figure 15:** TBE Maps During Hand, Tongue, & Foot Movement for Subject 19 (Page S16)

**Figure 16:** TBE Maps During Hand, Tongue, & Foot Movement for Subject 20 (Page S17)

**Reproduction of Speech Maps Across Subjects Not Shown in Main Article**

**Figure 17:** TBE Maps During Noun Production & Verb Generation Tasks for Subject 4 (Page S18)

**Figure 18:** TBE Maps During Noun Production & Verb Generation Tasks for Subject 5 (Page S19)

**Figure 19:** TBE Maps During Noun Production & Verb Generation Tasks for Subject 10 (Page S20)

**Figure 20:** TBE Maps During Verb Generation for Subject 11 (Page S21)

**Figure 21:** TBE Maps During Noun Production & Verb Generation Tasks for Subject 16 (Page S22)

**Figure 22:** TBE Maps During Noun Production & Verb Generation Tasks for Subject 17 (Page S23)

**Figure 23:** TBE Maps During Noun Production & Verb Generation Tasks for Subject 18 (Page S24)

**Figure 24:** TBE Maps During Noun Production & Verb Generation Tasks for Subject 19 (Page S25)

**Additional Results: White Matter Activity During Movement and Speech**

**Figure 25:** White Matter Activity During Hand, Tongue, & Foot Movement Across Subjects (Page S26)

**Figure 26:** White Matter Activity During Noun Production & Verb Generation Across Subjects (Page S27)

**Additional Results: Receiver Operating Characteristic (ROC) Curves Comparing Stimulation of Functional Mapping**

**Figure 27:** ROC Curves Comparing Stimulation and Functional Mapping of Speech Across All Subjects (Page S28)

**Figure 28:** ROC Curves Comparing Stimulation and Functional Mapping of Movement Across All Subjects (Page S29)

**Figure 29:** Potential Mechanisms Underlying Discordance Between Stimulation and Functional Mapping (Page S30)

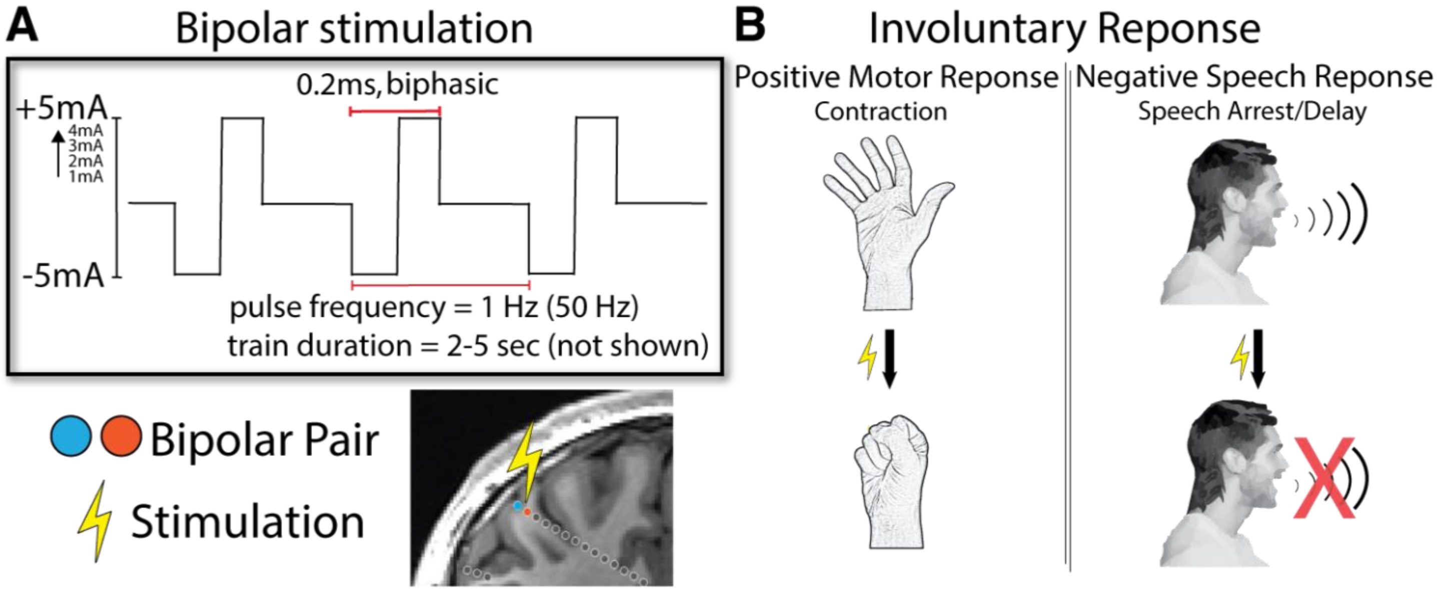


**Supplementary Figure 1. Clinical stimulation mapping using sEEG. A.** Bipolar stimulation between adjacent electrode contacts is delivered at cortical sites within and around regions of interest based on seizure patterns recorded during inpatient monitoring. **B.** Responses are observed by the clinical team or in the case of sensory phenomena (e.g. tingling), are reported by the patient. These evoked responses span many neurologic systems, but we limited our analyses to speech and sensorimotor related responses.

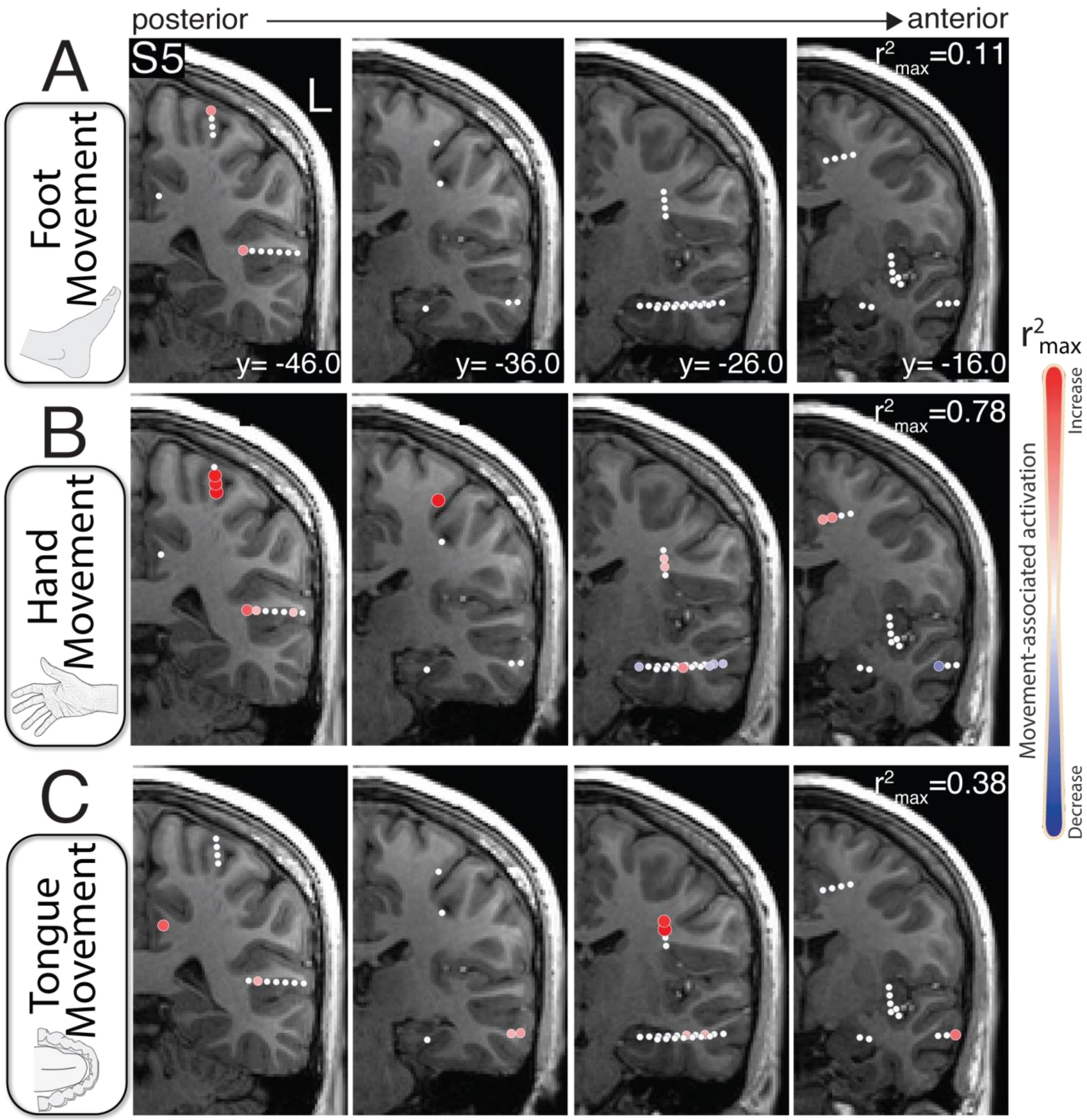


**Supplementary Figure 2. TBE maps during foot, hand and tongue movement – subject 5. A.** Coronal T1 slices showing channels (circles) that represent active (red) and inactive (white) tissue during foot movement as measured by movement specific broadband power shifts. **B.** As in A, but during hand movement. **C.** As in A, but during tongue movement.

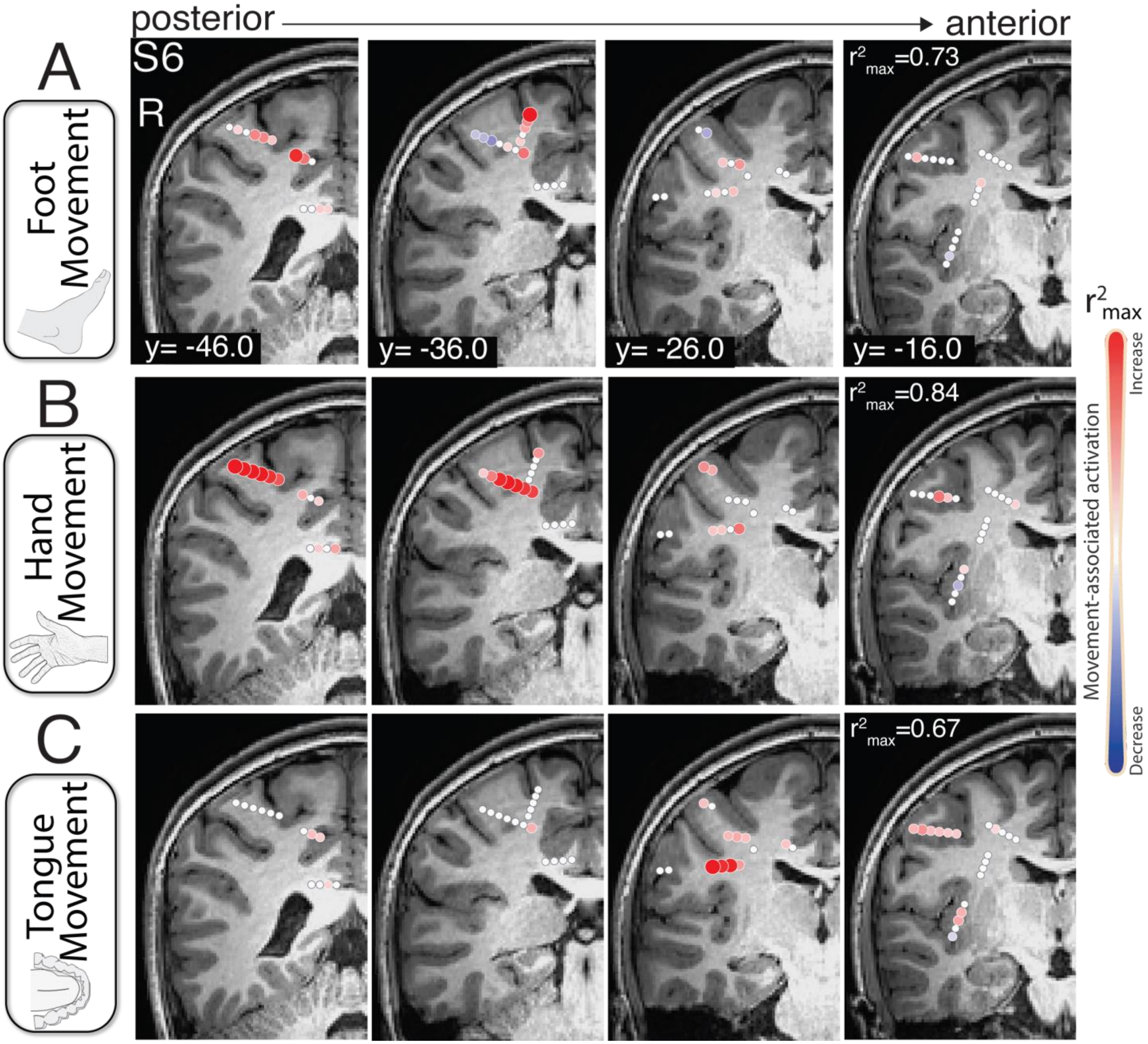


**Supplementary Figure 3. TBE maps during foot, hand and tongue movement – subject 6. A.** Coronal T1 slices showing channels (circles) that represent active (red) and inactive (white) tissue during foot movement as measured by movement specific broadband power shifts. **B.** As in A, but during hand movement. **C.** As in A, but during tongue movement.

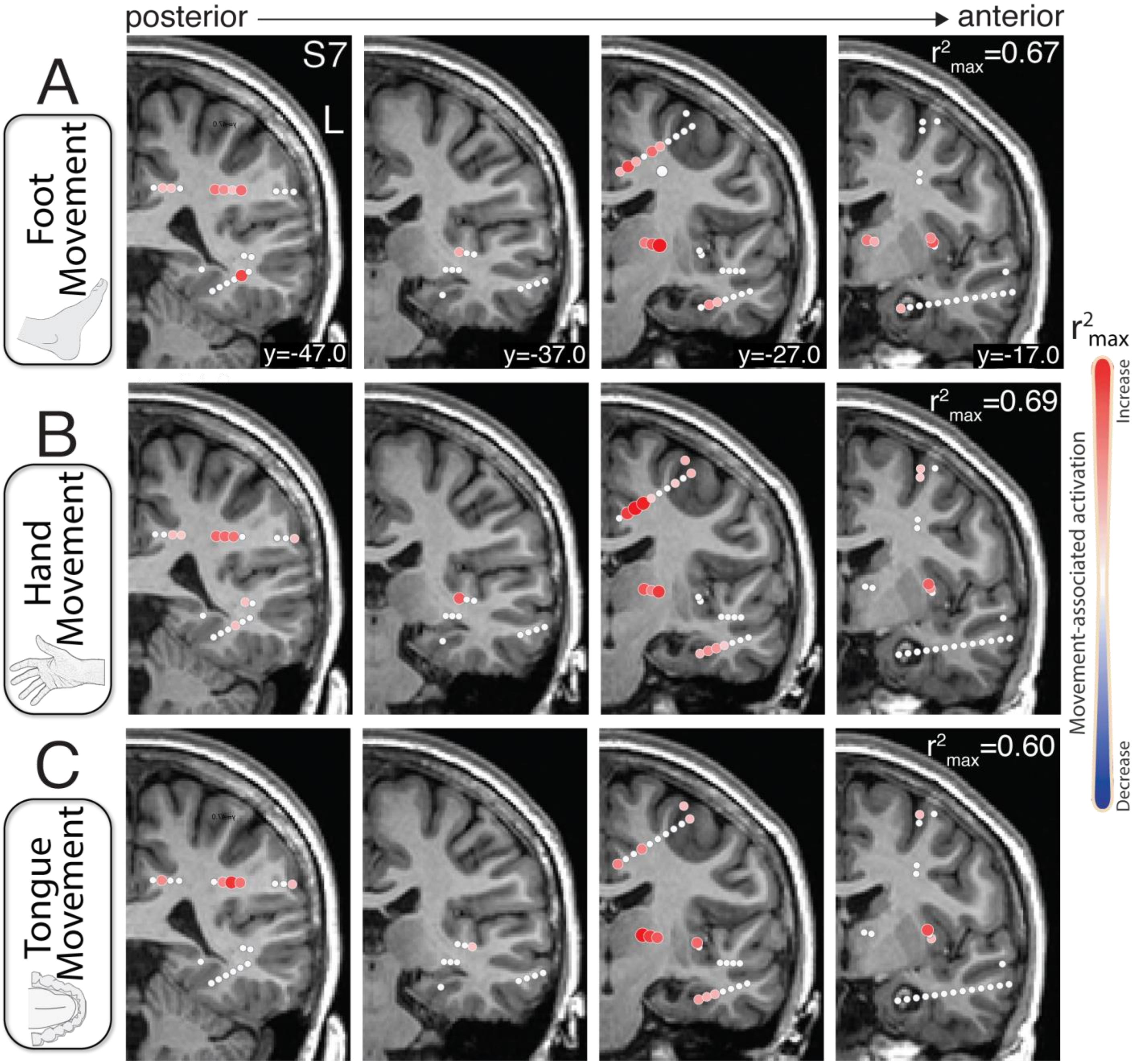


**Supplementary Figure 4. TBE maps during foot, hand and tongue movement – subject 7. A.** Coronal T1 slices showing channels (circles) that represent active (red) and inactive (white) tissue during foot movement as measured by movement specific broadband power shifts. **B.** As in A, but during hand movement. **C.** As in A, but during tongue movement.

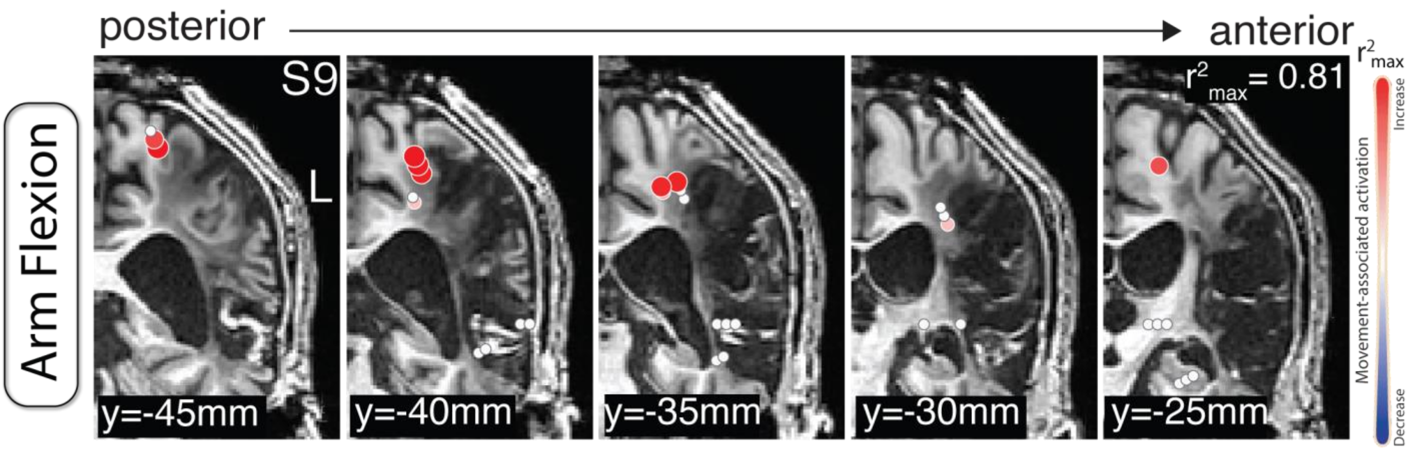


**Supplementary Figure 5. TBE maps during arm flexion – subject 9.** Coronal T1 slices showing channels (circles) that represent active (red) and inactive (white) tissue during arm flexion as measured by movement specific broadband power shifts.

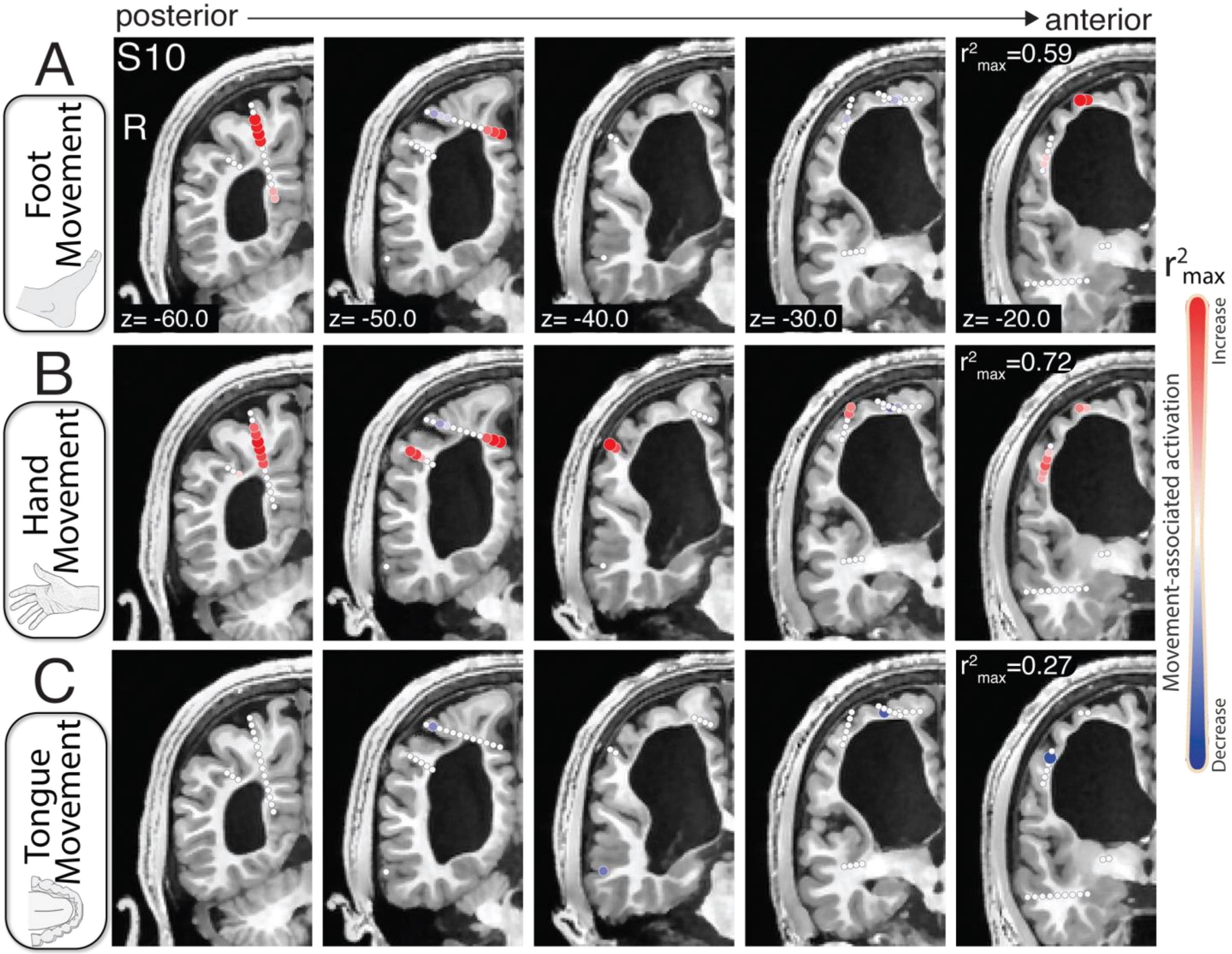


**Supplementary Figure 6. TBE maps during foot, hand and tongue movement – subject 10. A.** Coronal T1 slices showing channels (circles) that represent active (red) and inactive (white) tissue during foot movement as measured by movement specific broadband power shifts. **B.** As in A, but during hand movement. **C.** As in A, but during tongue movement.

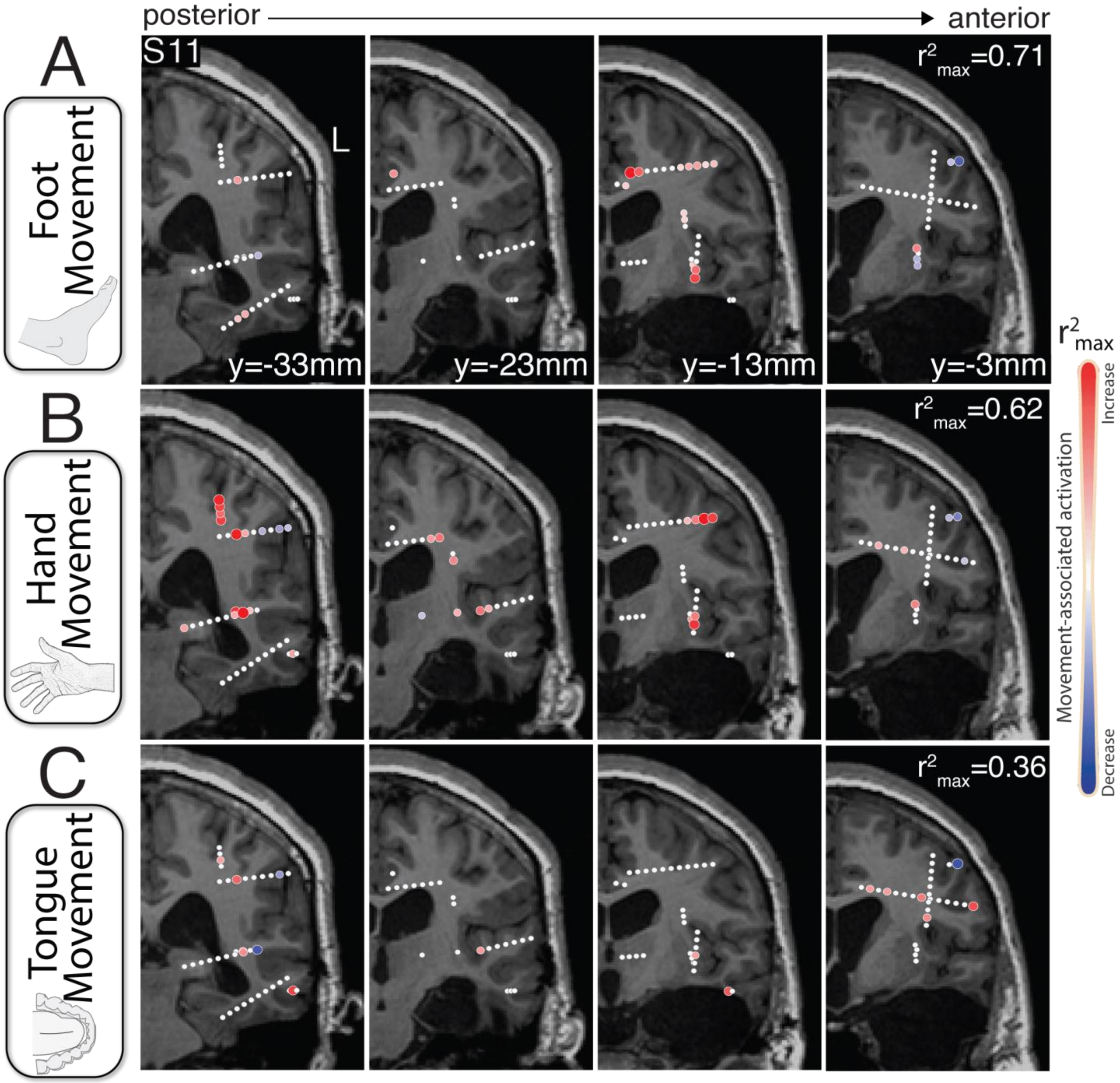


**Supplementary Figure 7. TBE maps during foot, hand and tongue movement – subject 11. A.** Coronal T1 slices showing channels (circles) that represent active (red) and inactive (white) tissue during foot movement as measured by movement specific broadband power shifts. **B.** As in A, but during hand movement. **C.** As in A, but during tongue movement.

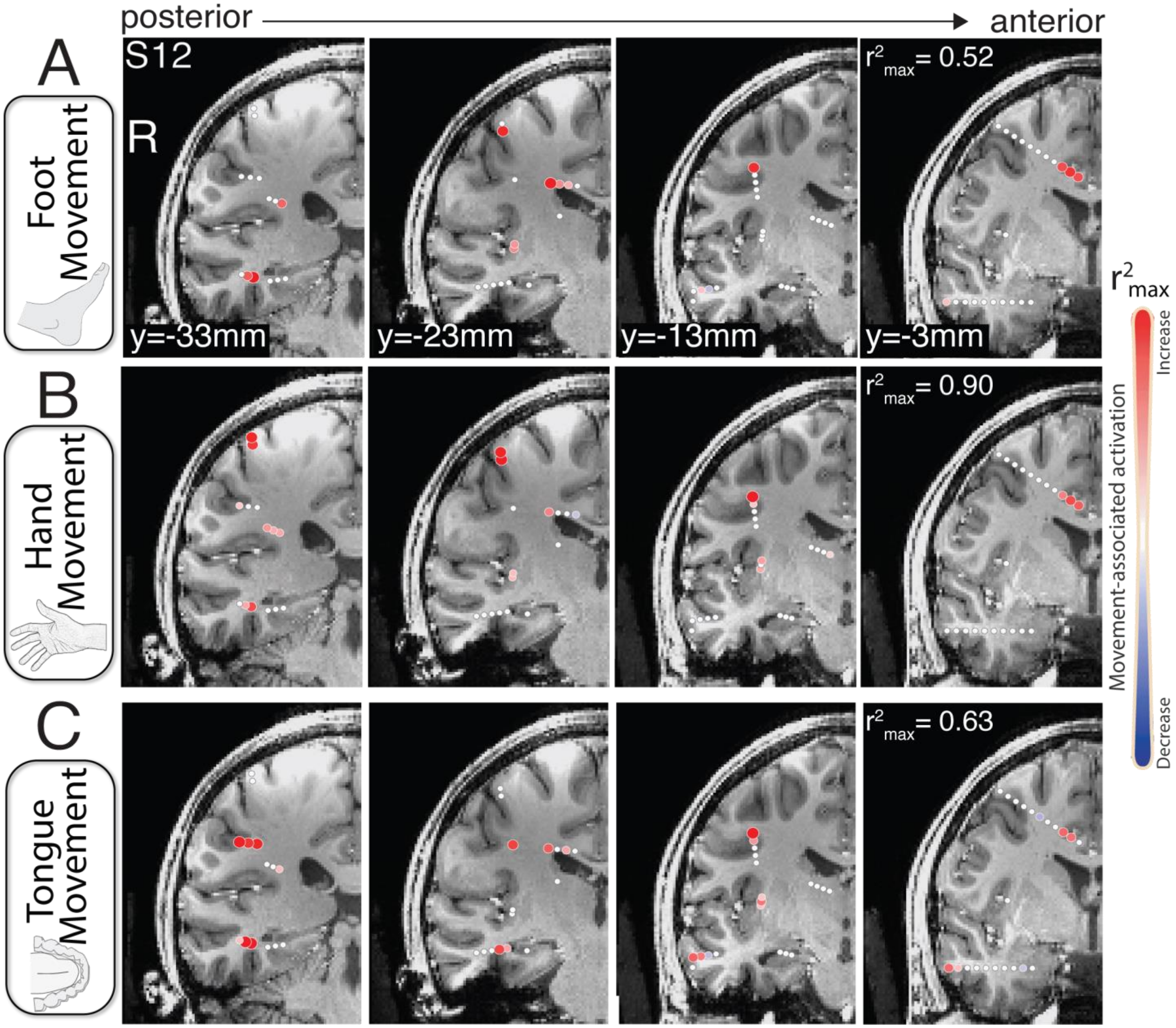


**Supplementary Figure 8. TBE maps during foot, hand and tongue movement – subject 12. A.** Coronal T1 slices showing channels (circles) that represent active (red) and inactive (white) tissue during foot movement as measured by movement specific broadband power shifts. **B.** As in A, but during hand movement. **C.** As in A, but during tongue movement.

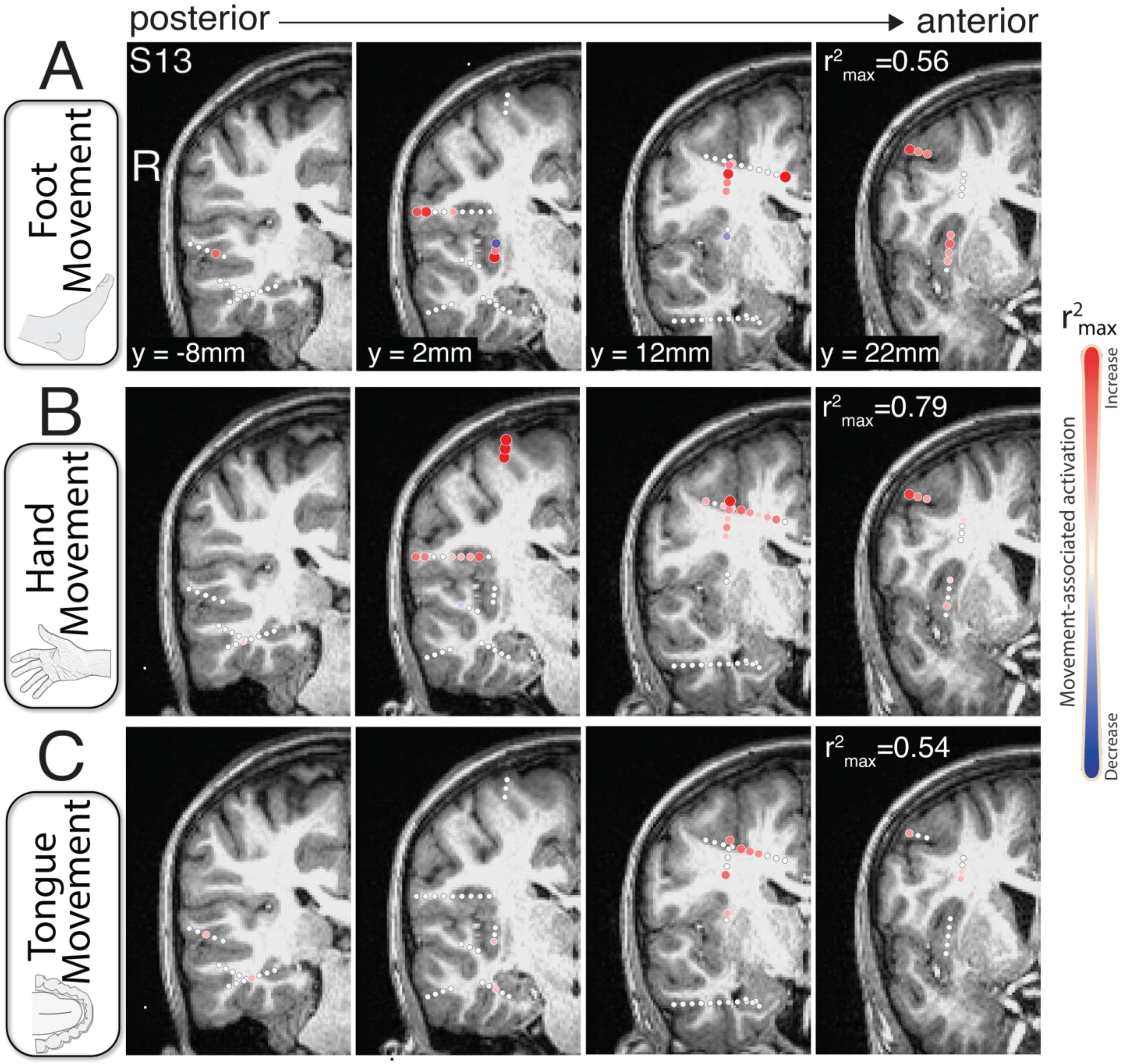


**Supplementary Figure 9. TBE maps during foot, hand and tongue movement – subject 13. A.** Coronal T1 slices showing channels (circles) that represent active (red) and inactive (white) tissue during foot movement as measured by movement specific broadband power shifts. **B.** As in A, but during hand movement. **C.** As in A, but during tongue movement.

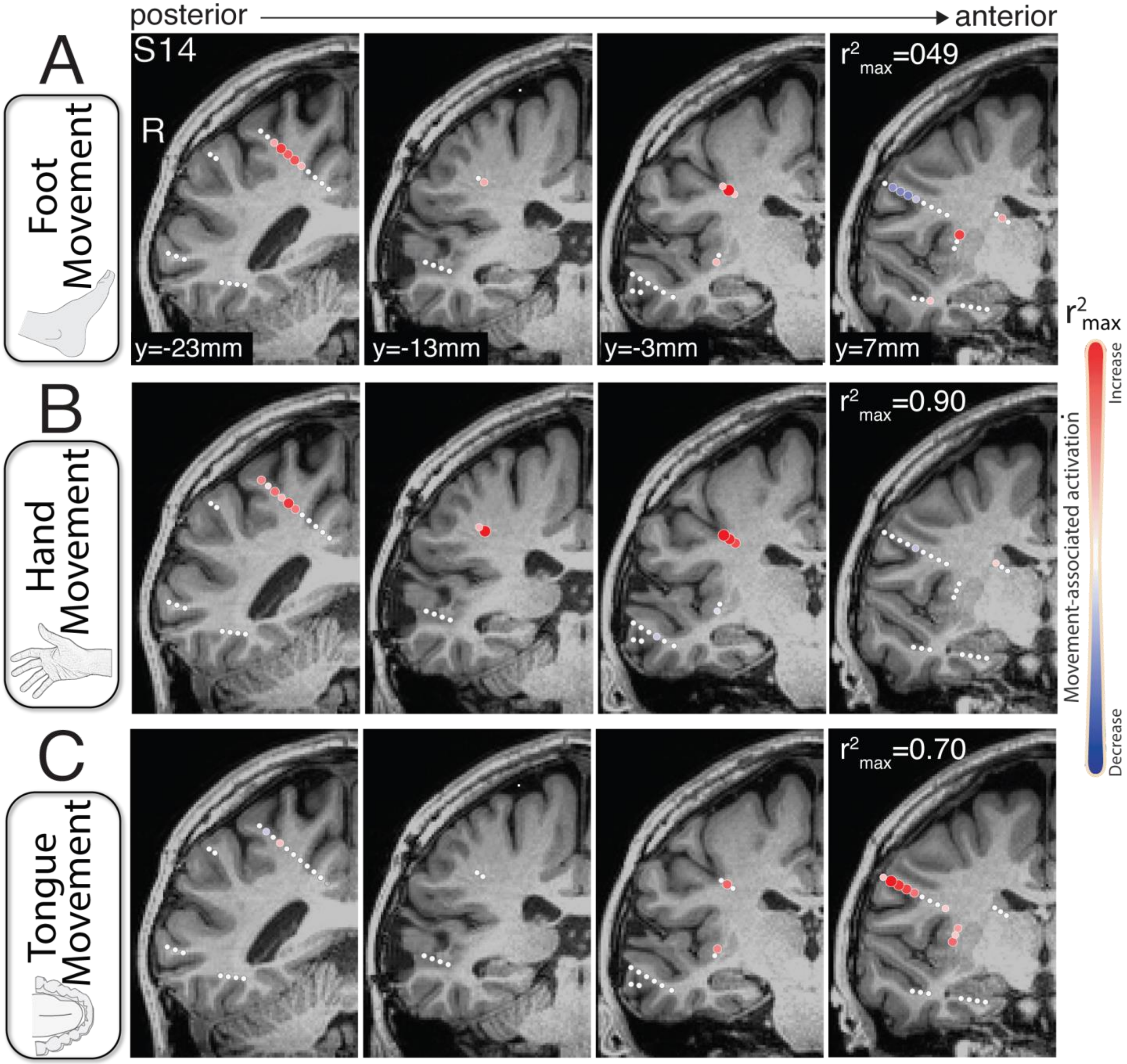


**Supplementary Figure 10. TBE maps during foot, hand and tongue movement – subject 14. A.** Coronal T1 slices showing channels (circles) that represent active (red) and inactive (white) tissue during foot movement as measured by movement specific broadband power shifts. **B.** As in A, but during hand movement. **C.** As in A, but during tongue movement.

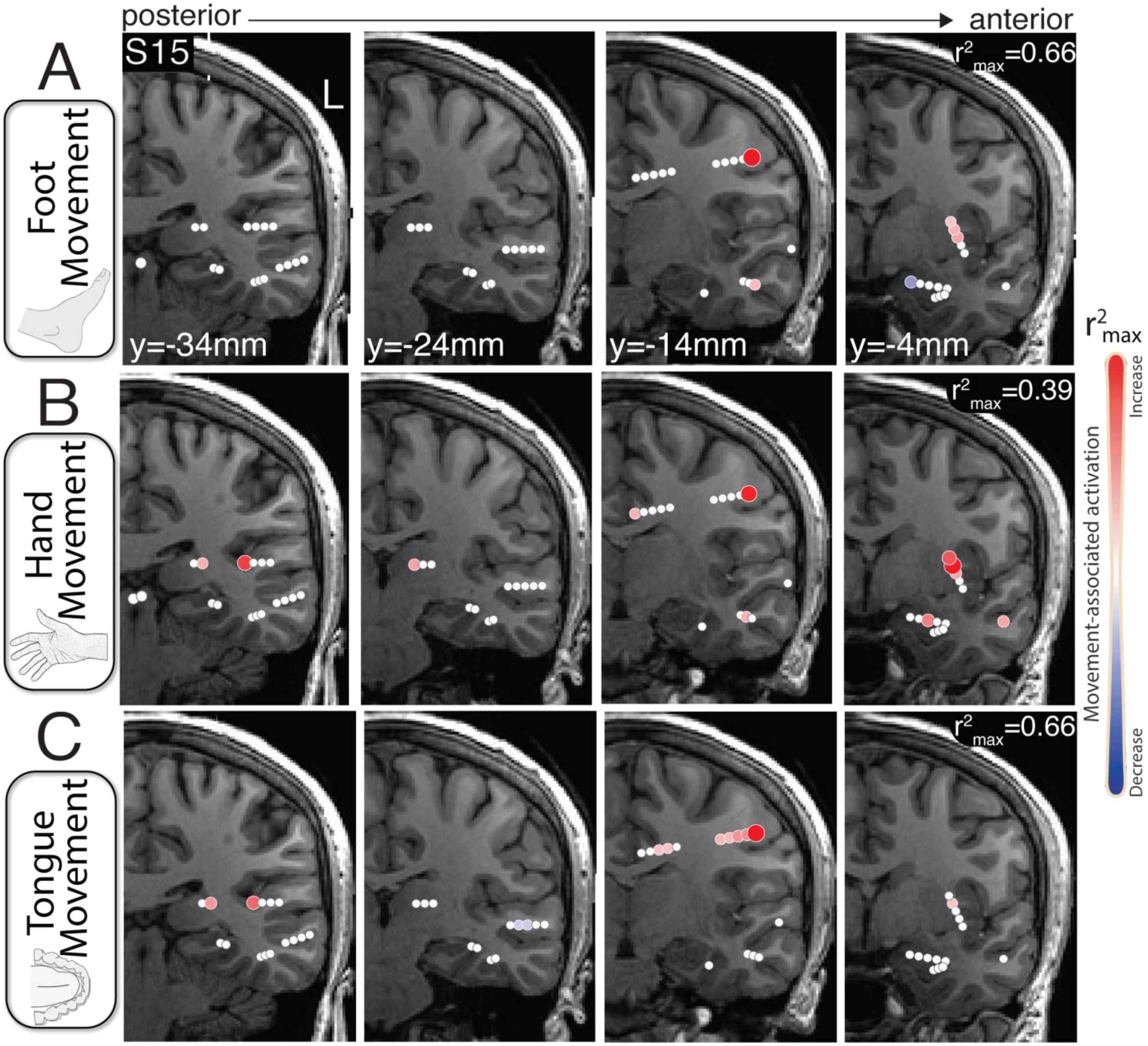


**Supplementary Figure 11. TBE maps during foot, hand and tongue movement – subject 15. A.** Coronal T1 slices showing channels (circles) that represent active (red) and inactive (white) tissue during foot movement as measured by movement specific broadband power shifts. **B.** As in A, but during hand movement. **C.** As in A, but during tongue movement.

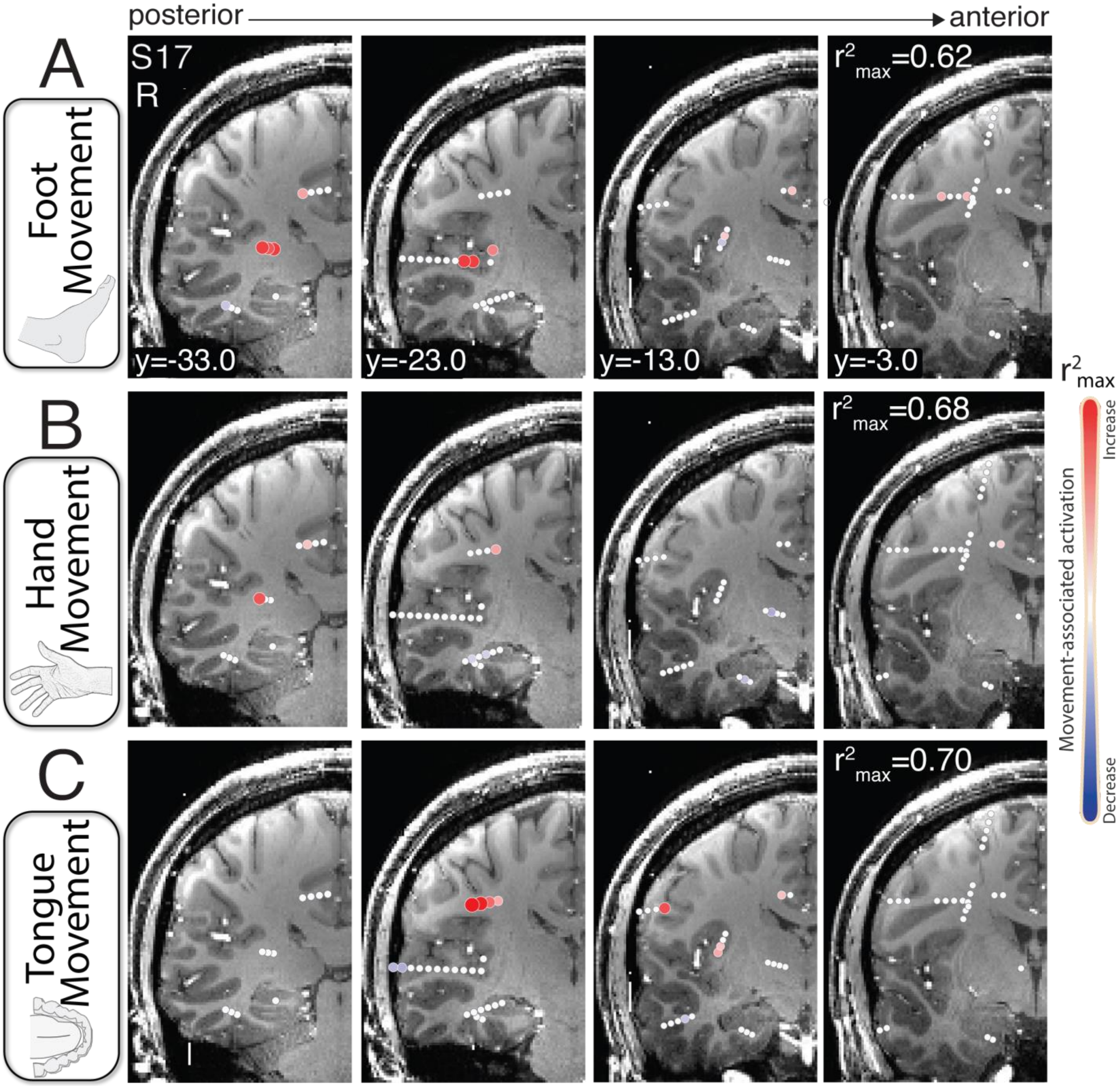


**Supplementary Figure 12. TBE maps during foot, hand and tongue movement – subject 17. A.** Coronal T1 slices showing channels (circles) that represent active (red) and inactive (white) tissue during foot movement as measured by movement specific broadband power shifts. **B.** As in A, but during hand movement. **C.** As in A, but during tongue movement.

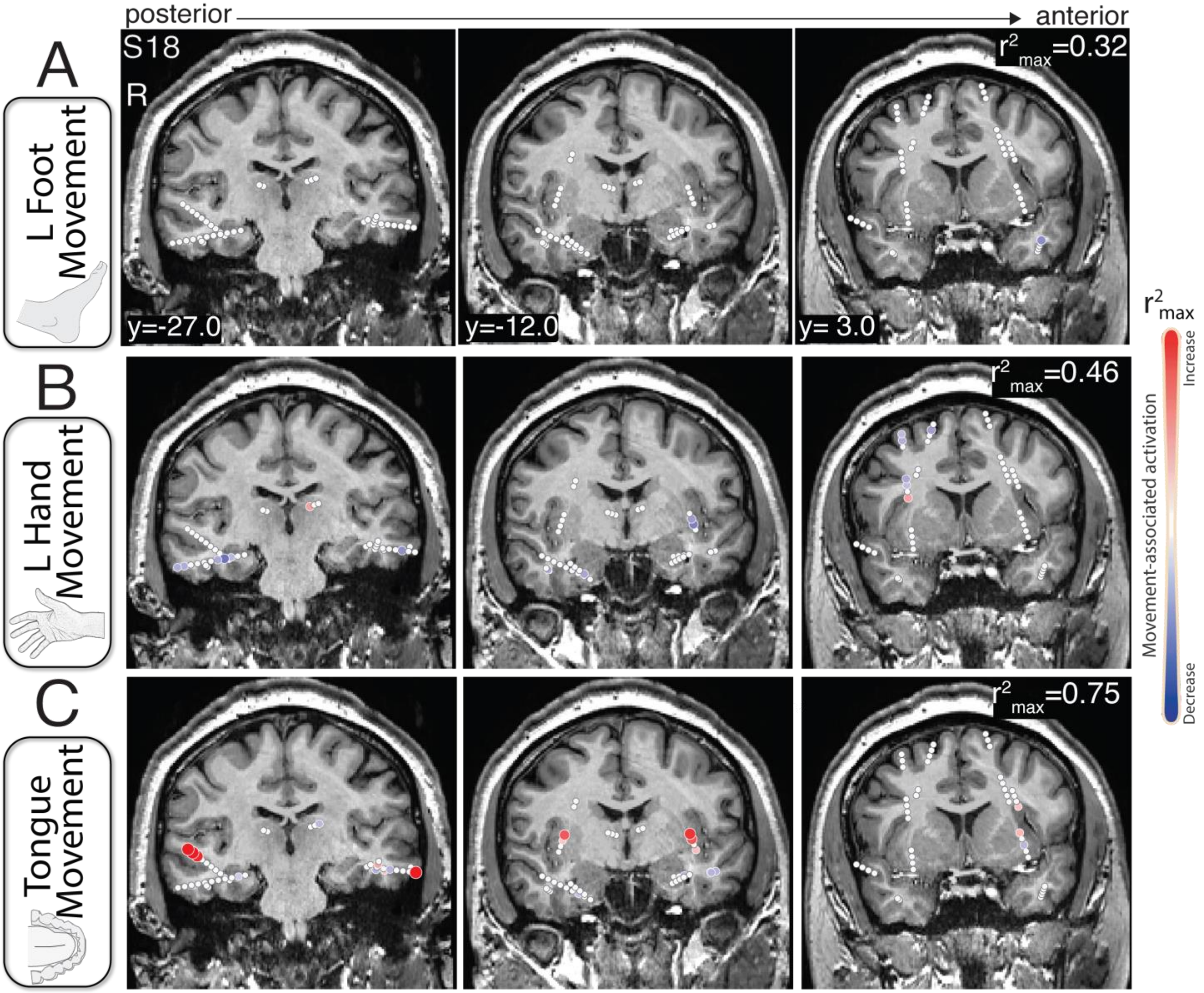


**Supplementary Figure 13. TBE maps during left foot, hand and tongue movement – subject 18. A.** Coronal T1 slices showing channels (circles) that represent active (red) and inactive (white) tissue during foot movement as measured by movement specific broadband power shifts. **B.** As in A, but during hand movement. **C.** As in A, but during tongue movement.

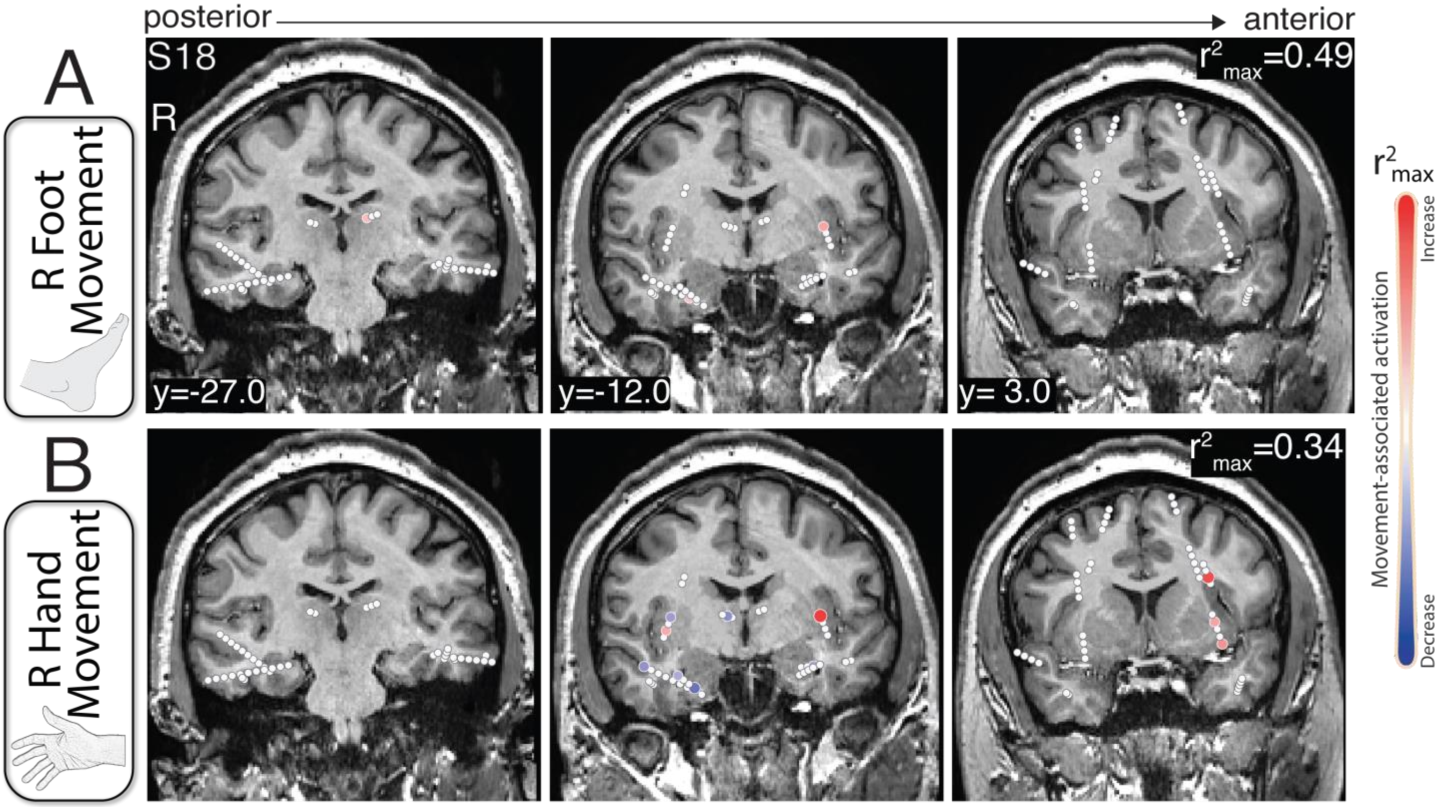


**Supplementary Figure 14. TBE maps during right foot and hand movement – subject 18. A.** Coronal T1 slices showing channels (circles) that represent active (red) and inactive (white) tissue during foot movement as measured by movement specific broadband power shifts. **B.** As in A, but during hand movement.

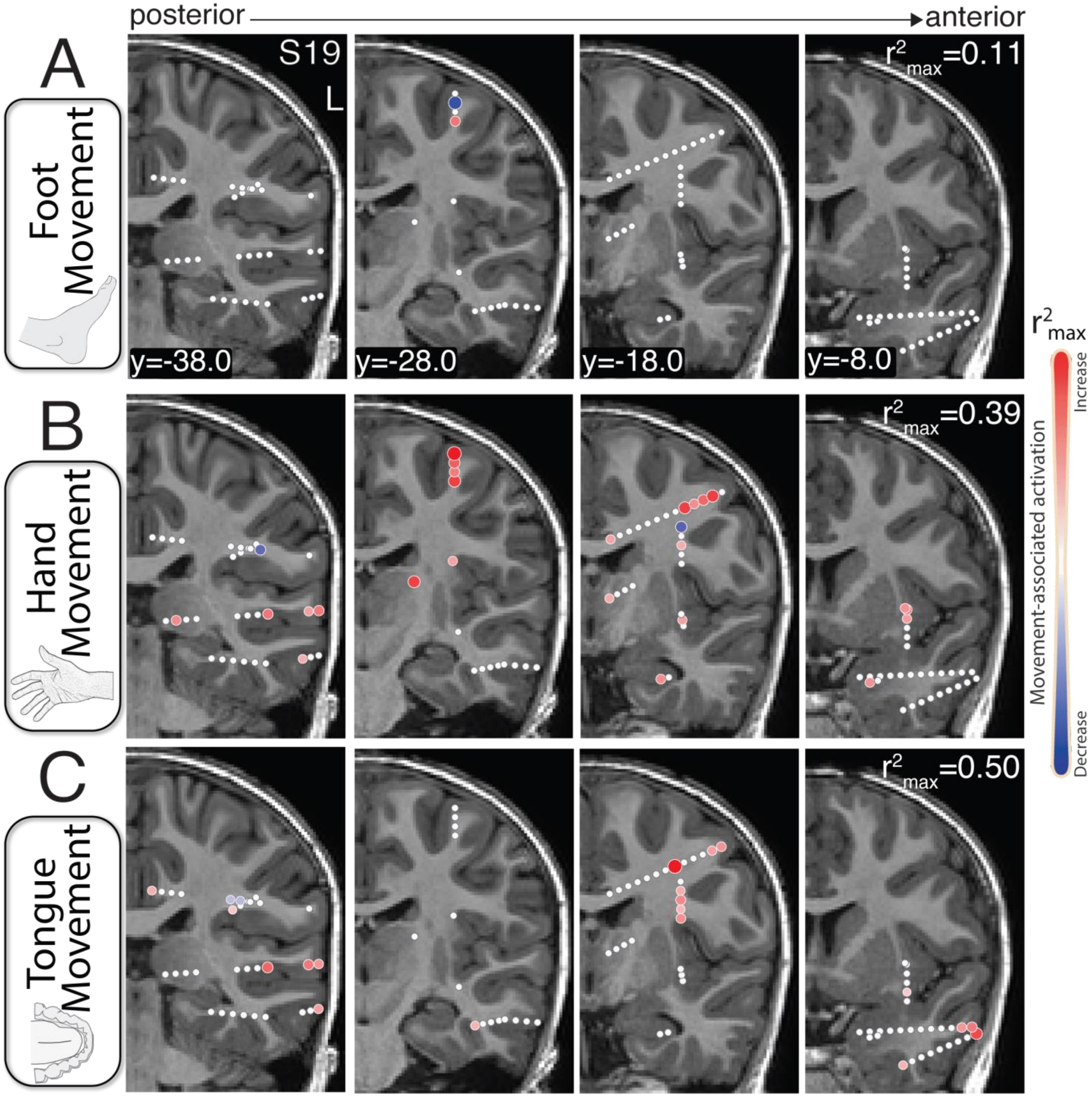


**Supplementary Figure 15. TBE maps during foot, hand and tongue movement – subject 19. A.** Coronal T1 slices showing channels (circles) that represent active (red) and inactive (white) tissue during foot movement as measured by movement specific broadband power shifts. **B.** As in A, but during hand movement. **C.** As in A, but during tongue movement.

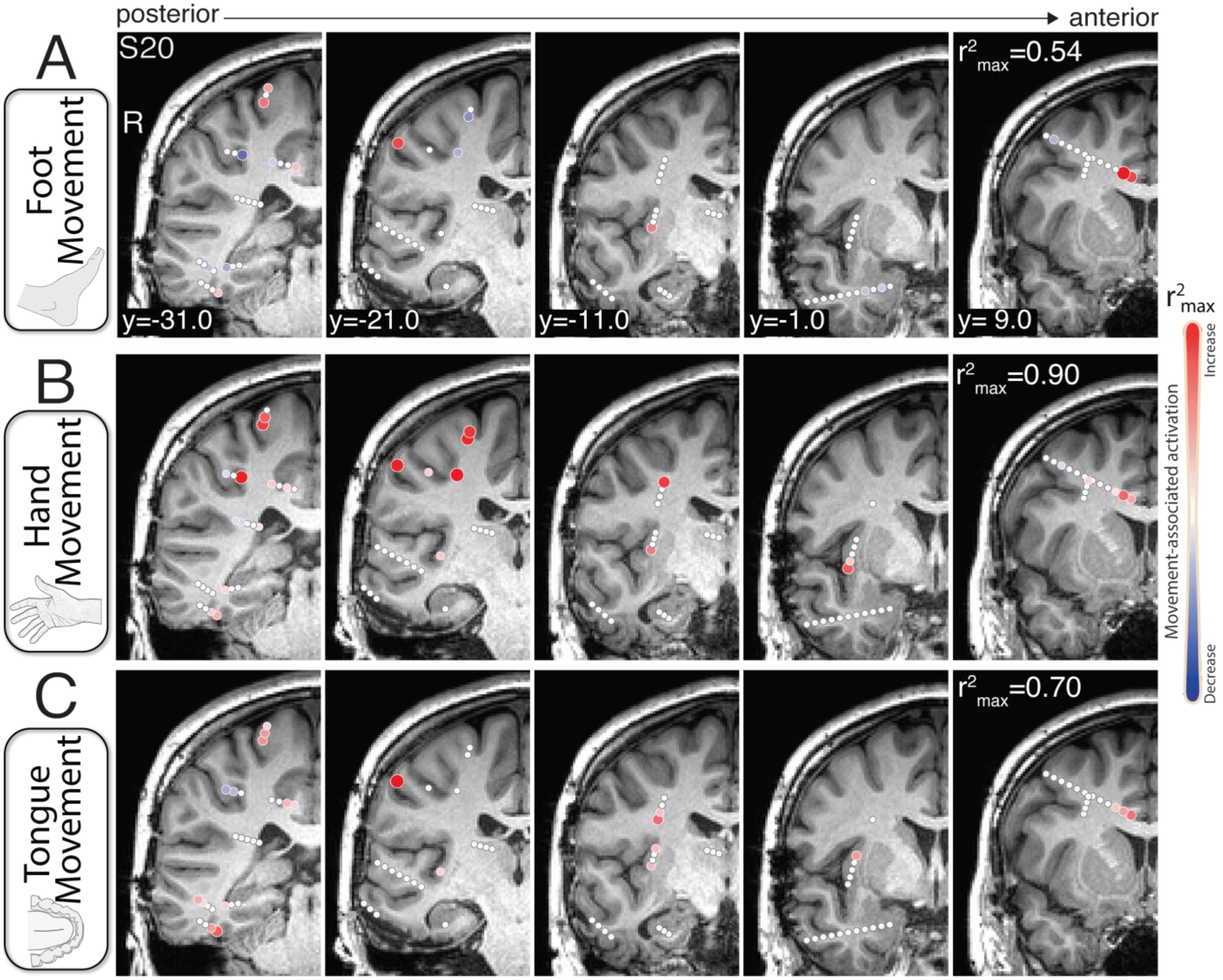


**Supplementary Figure 16. TBE maps during foot, hand and tongue movement – subject 20. A.** Coronal T1 slices showing channels (circles) that represent active (red) and inactive (white) tissue during foot movement as measured by movement specific broadband power shifts. **B.** As in A, but during hand movement. **C.** As in A, but during tongue movement.

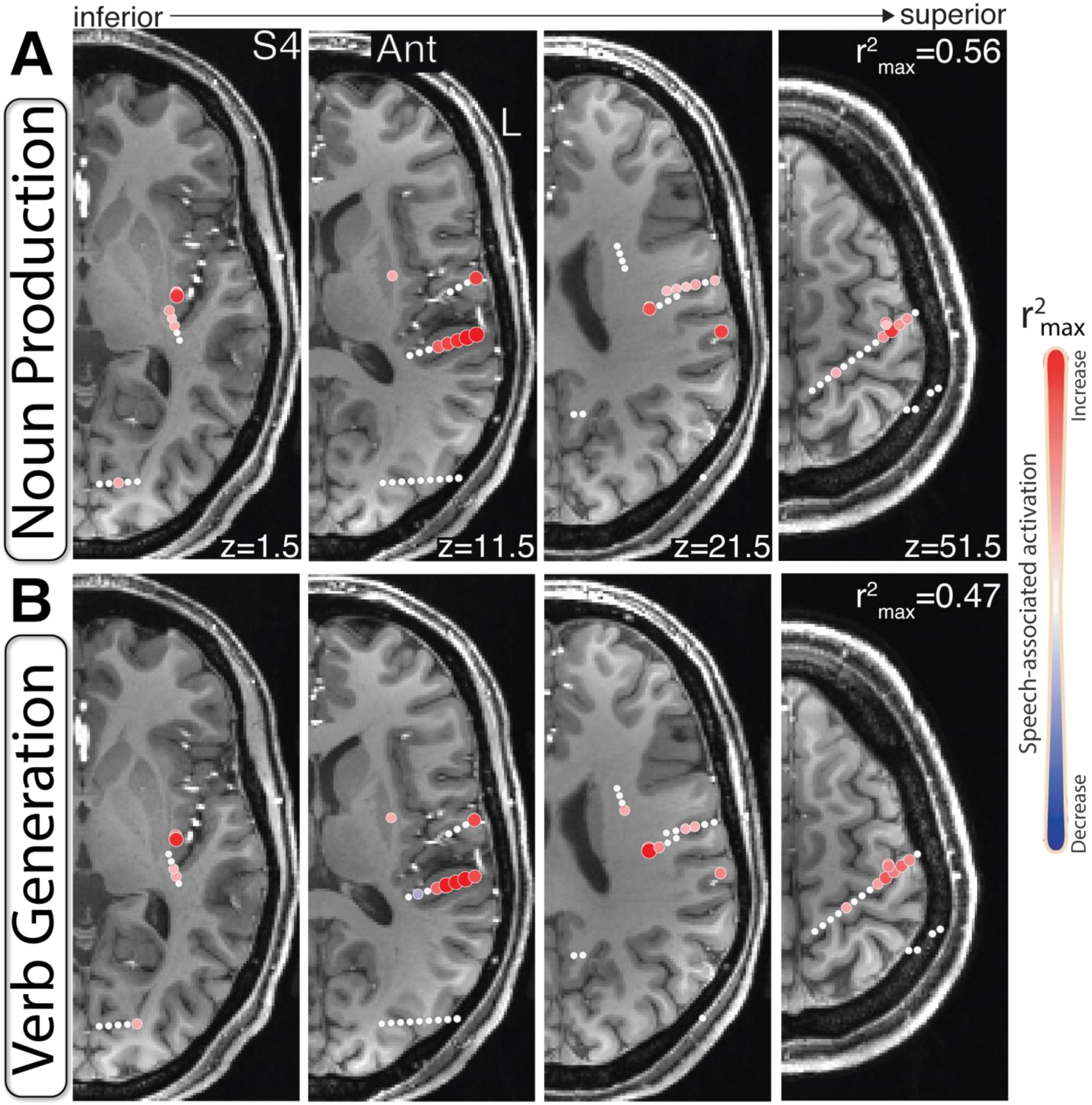


**Supplementary Figure 17. TBE maps during speech tasks – subject 4. A.** Axial T1 slices showing channels (circles) that represent active (red) and inactive (white) tissue during a noun production task as measured by movement specific broadband power shifts. **B.** As in A, but during a verb generation task.

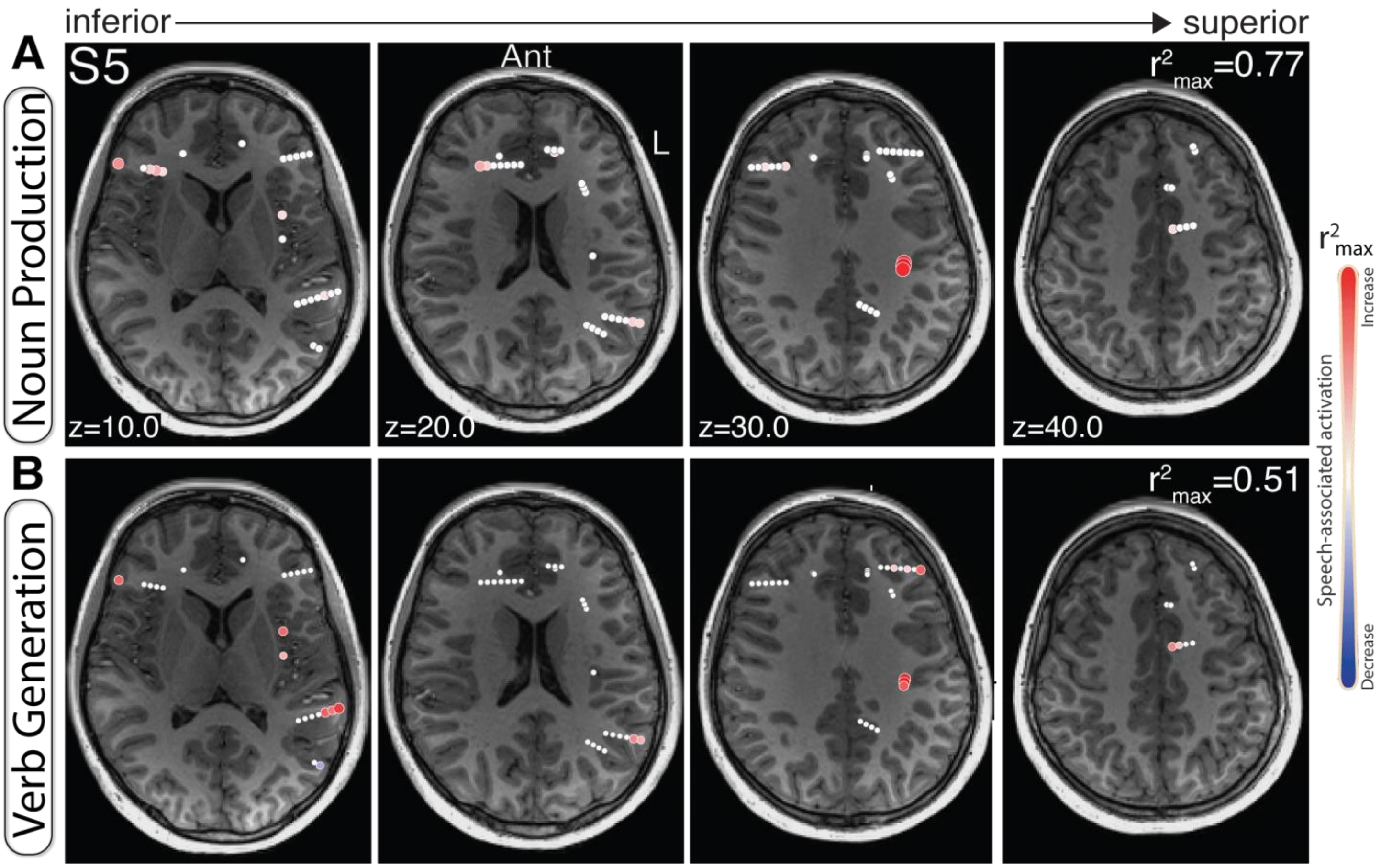


**Supplementary Figure 18. TBE maps during speech tasks – subject 5. A.** Axial T1 slices showing channels (circles) that represent active (red) and inactive (white) tissue during a noun production task as measured by movement specific broadband power shifts. **B.** As in A, but during a verb generation task.

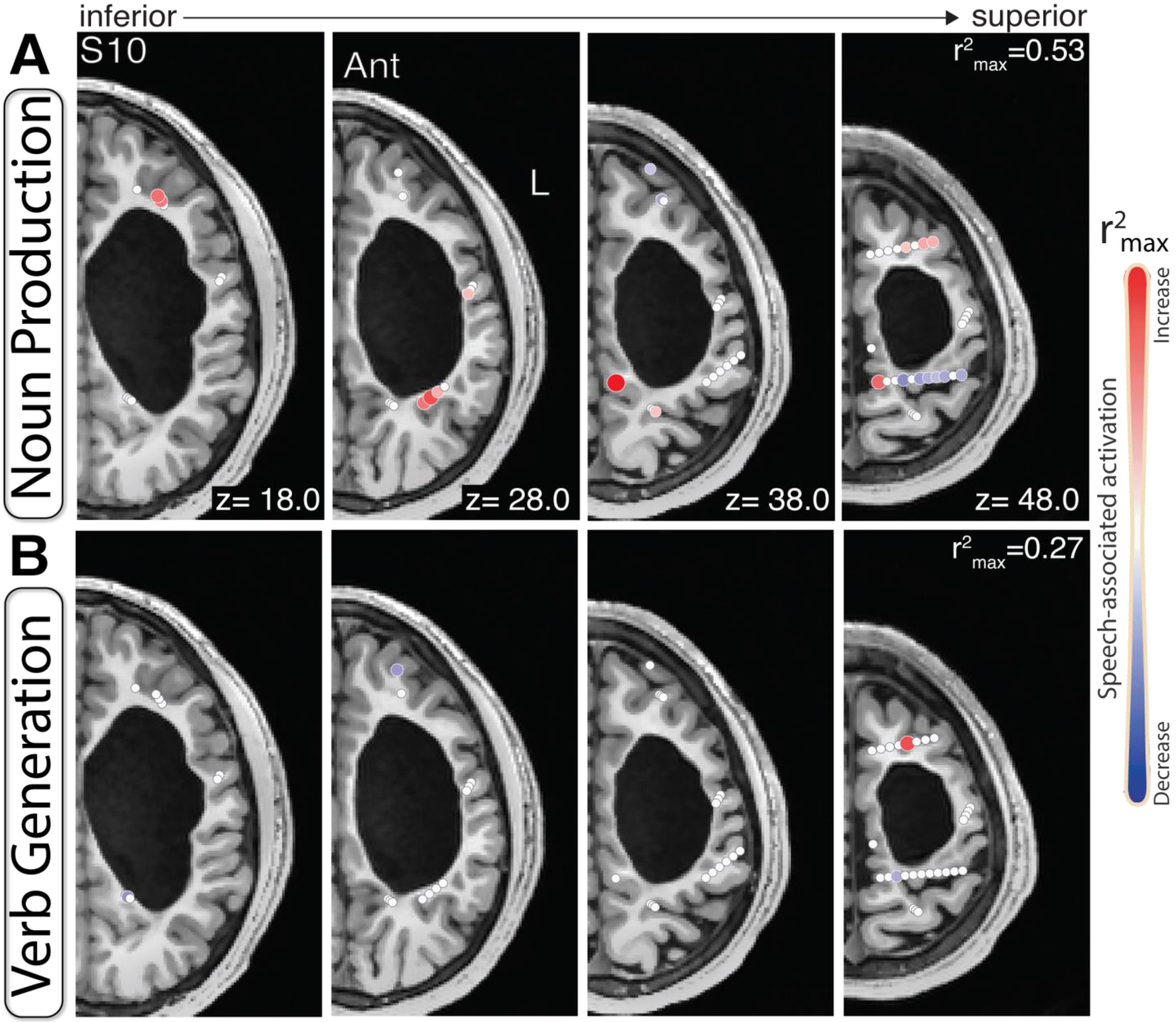


**Supplementary Figure 19. TBE maps during speech tasks – subject 10. A.** Axial T1 slices showing channels (circles) that represent active (red) and inactive (white) tissue during a noun production task as measured by movement specific broadband power shifts. **B.** As in A, but during a verb generation task.

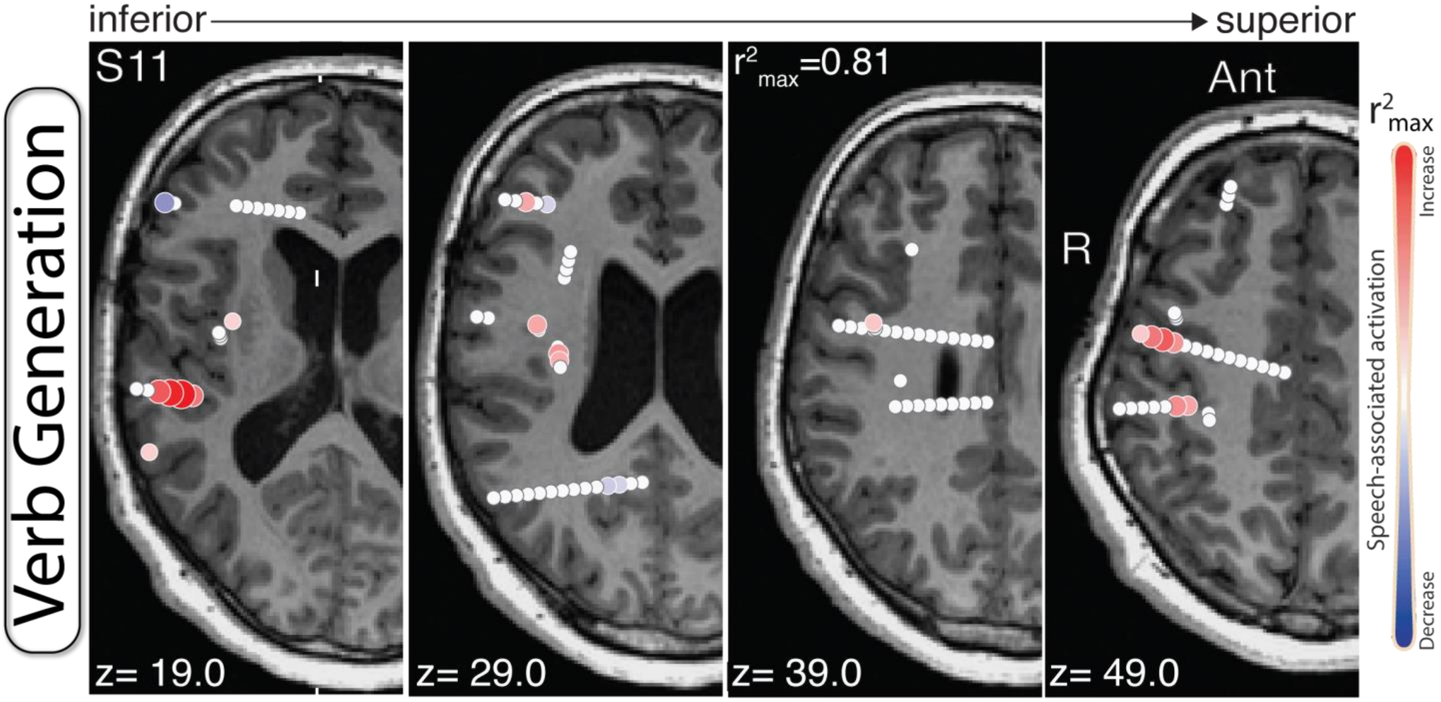


**Supplementary Figure 20. TBE maps during speech tasks – subject 11. A.** Axial T1 slices showing channels (circles) that represent active (red) and inactive (white) tissue during a verb generation task as measured by movement specific broadband power shifts. Note: Subject 7 was determined to be right language dominant by fMRI.

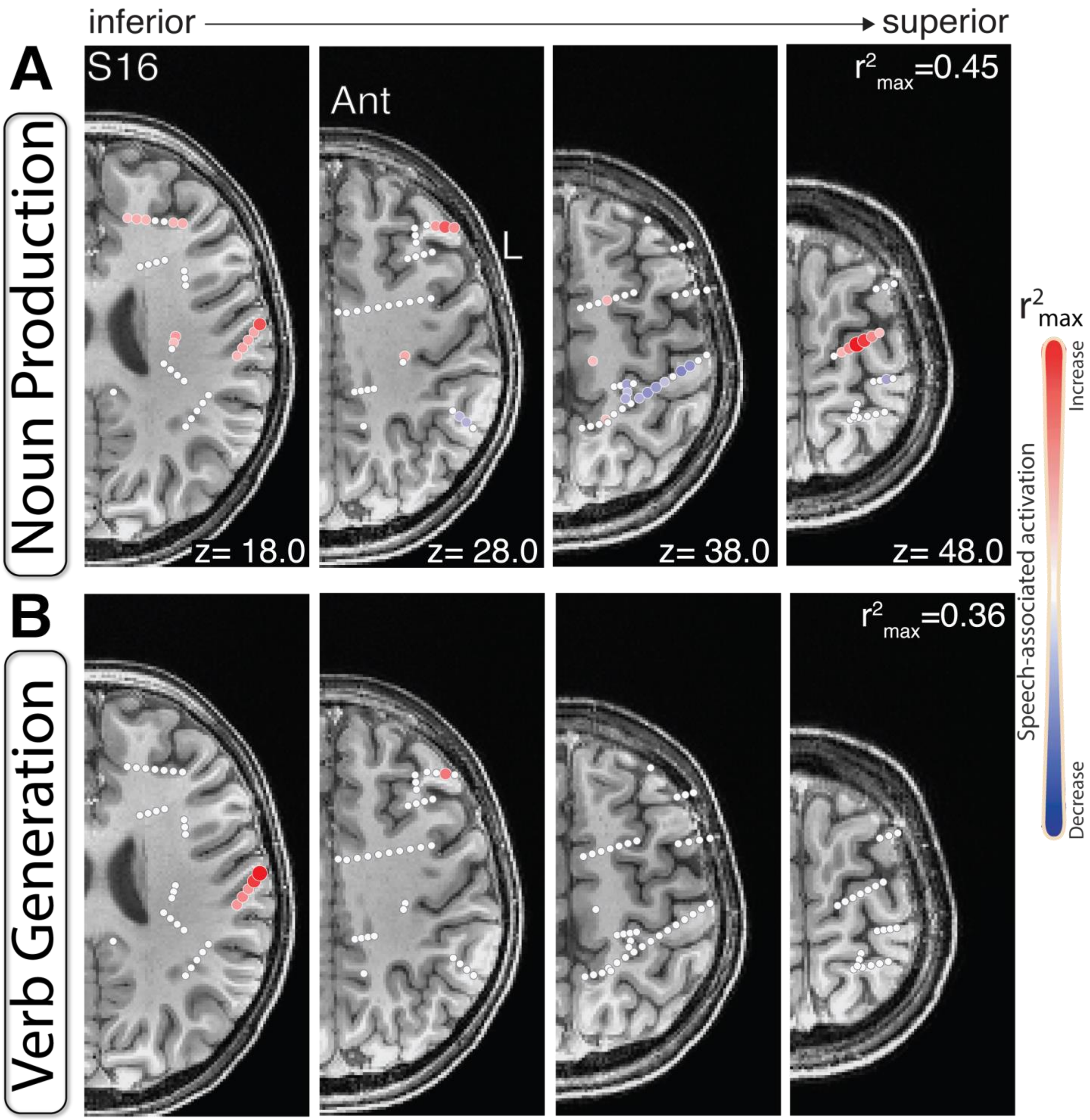


**Supplementary Figure 21. TBE maps during speech tasks – subject 16. A.** Axial T1 slices showing channels (circles) that represent active (red) and inactive (white) tissue during a noun production task as measured by movement specific broadband power shifts. **B.** As in A, but during a verb generation task. Note: No significant broadband shifts were measured during either task. Thus this demonstrated a negative result.

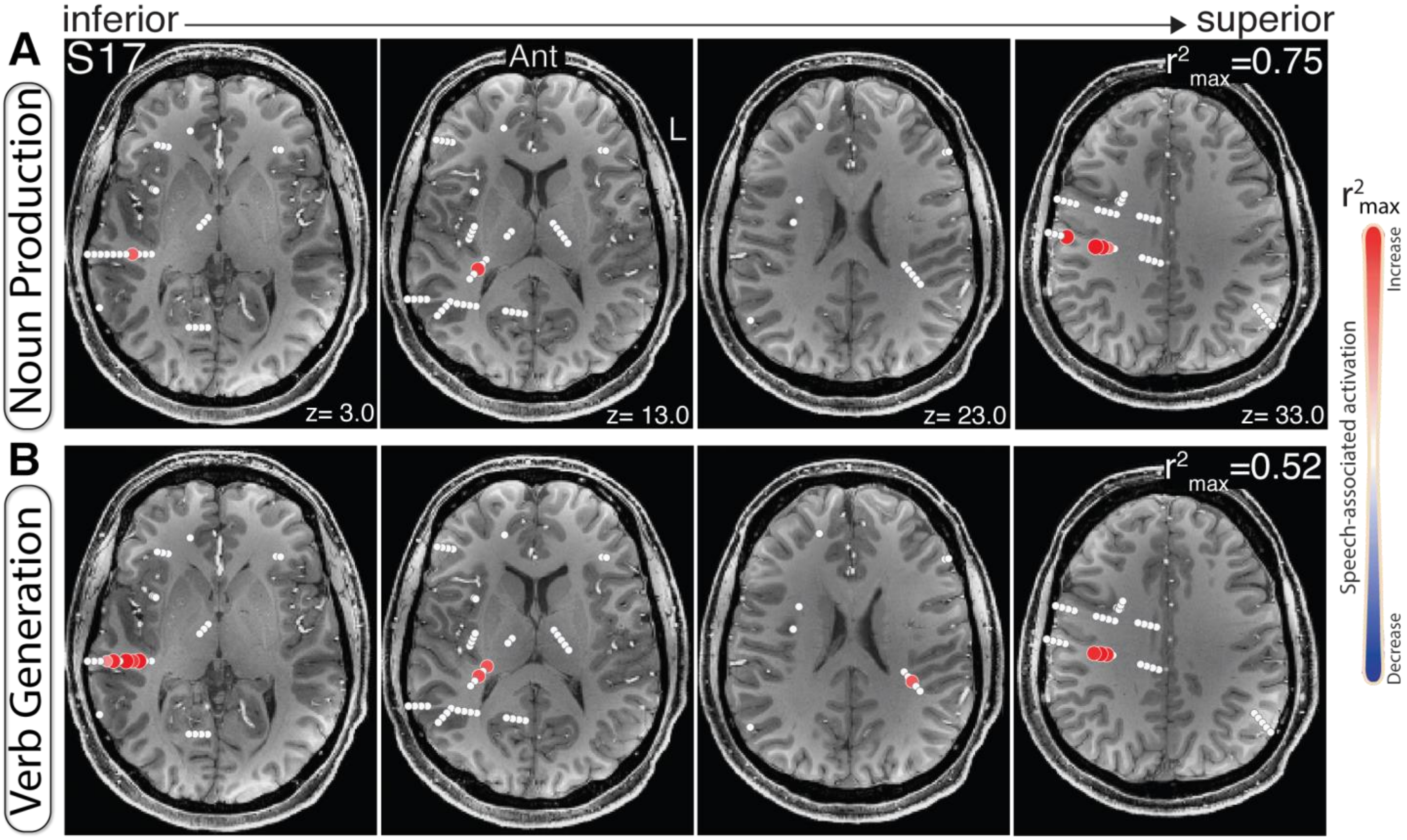


**Supplementary Figure 22. TBE maps during speech tasks– subject 17. A.** Axial T1 slices showing channels (circles) that represent active (red) and inactive (white) tissue during a noun production task as measured by movement specific broadband power shifts. **B.** As in A, but during a verb generation task.

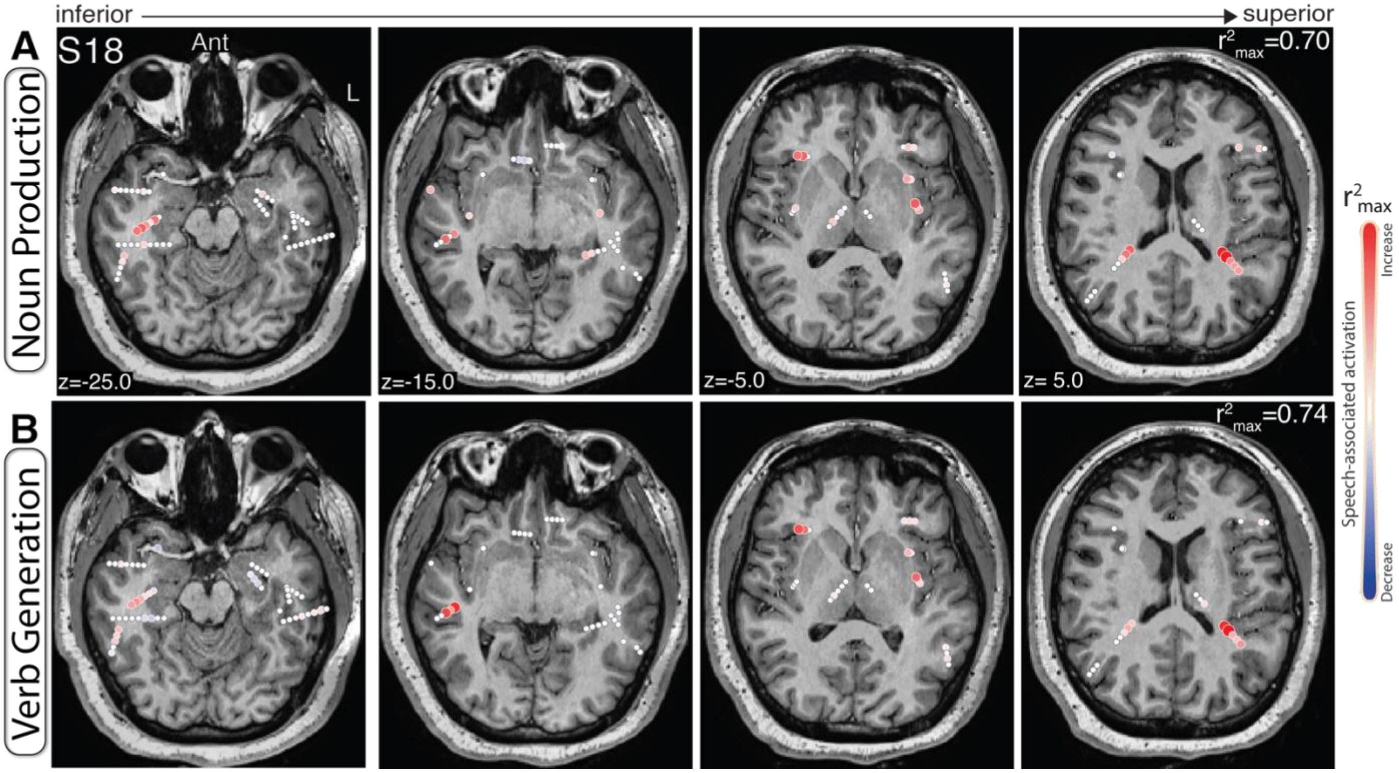


**Supplementary Figure 23. TBE maps during speech tasks– subject 18. A.** Axial T1 slices showing channels (circles) that represent active (red) and inactive (white) tissue during a noun production task as measured by movement specific broadband power shifts. **B.** As in A, but during a verb generation task.

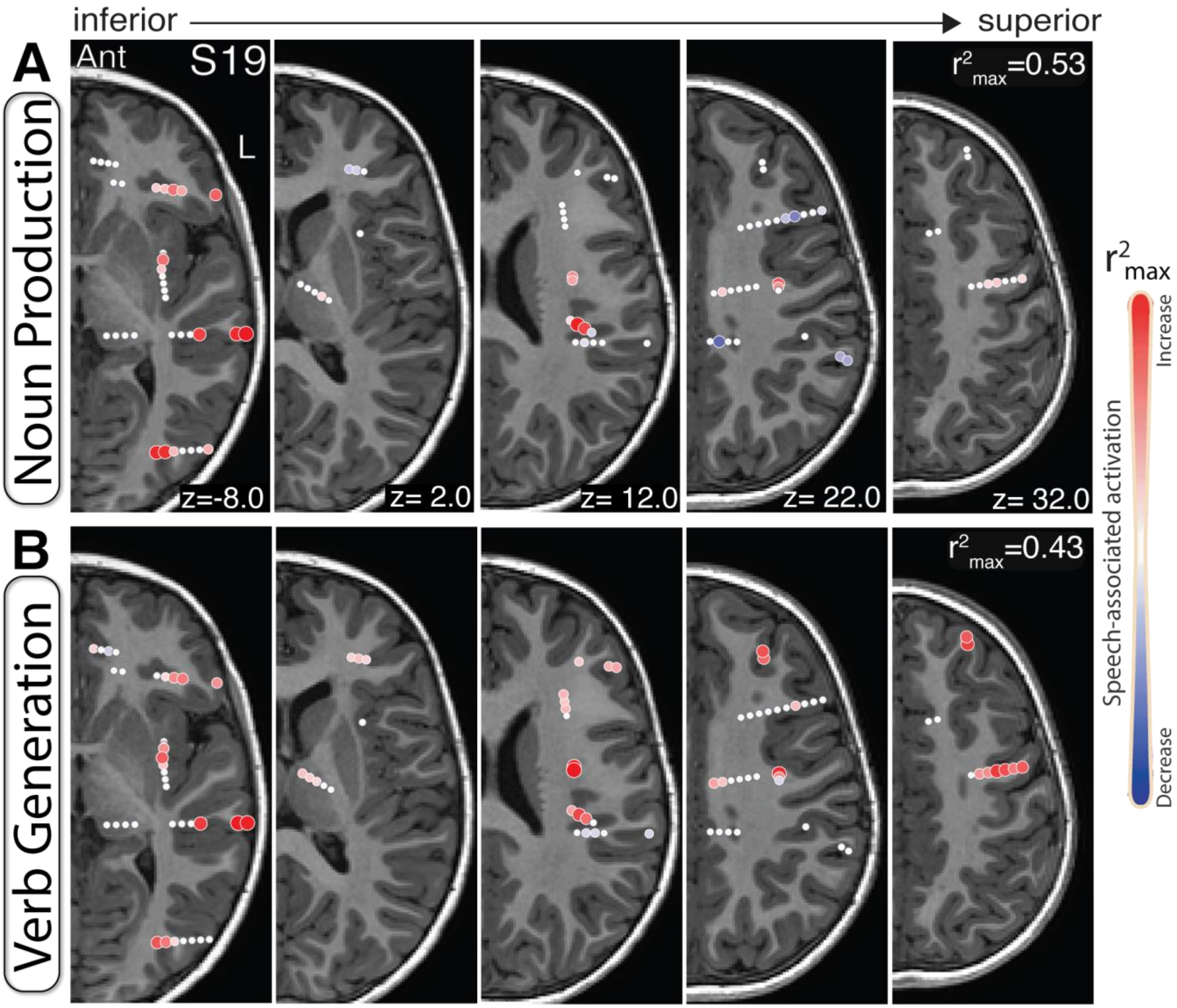


**Supplementary Figure 24. TBE maps during speech tasks– subject 19. A.** Axial T1 slices showing channels (circles) that represent active (red) and inactive (white) tissue during a noun production task as measured by movement specific broadband power shifts. **B.** As in A, but during a verb generation task.

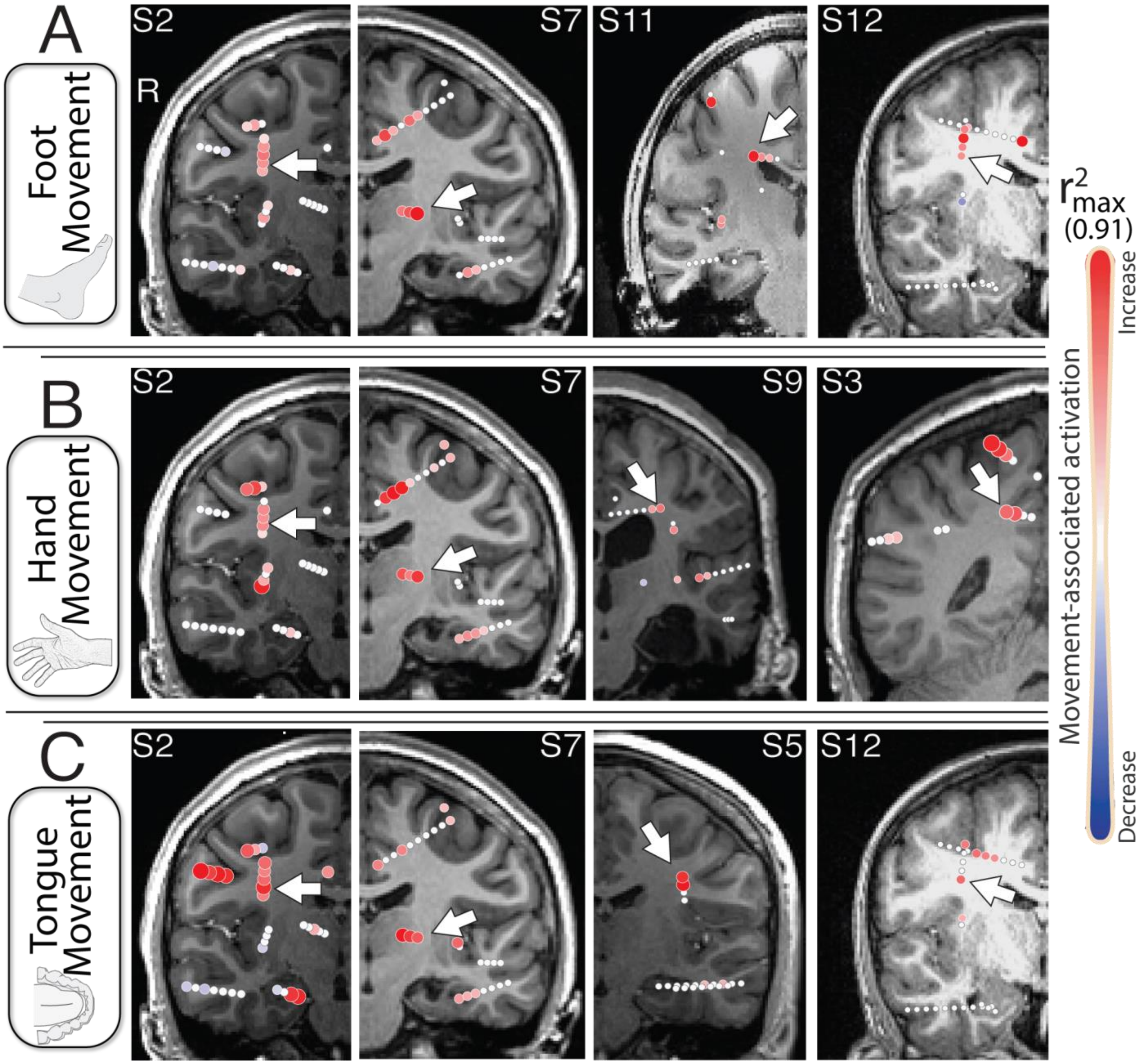


**Supplementary Figure 25. White matter activity during motor mapping. A.** T1 coronal slices showing activity maps demonstrating broadband power shifts in white matter (white arrow) during foot movement. **B.** As in A, but during hand movement. **C.** As in A, but during tongue movement. Note: The purpose of this figure is to illustrate significant activity in white matter. The $r^2$ values are not scaled to one another as this is not essential to the figure.

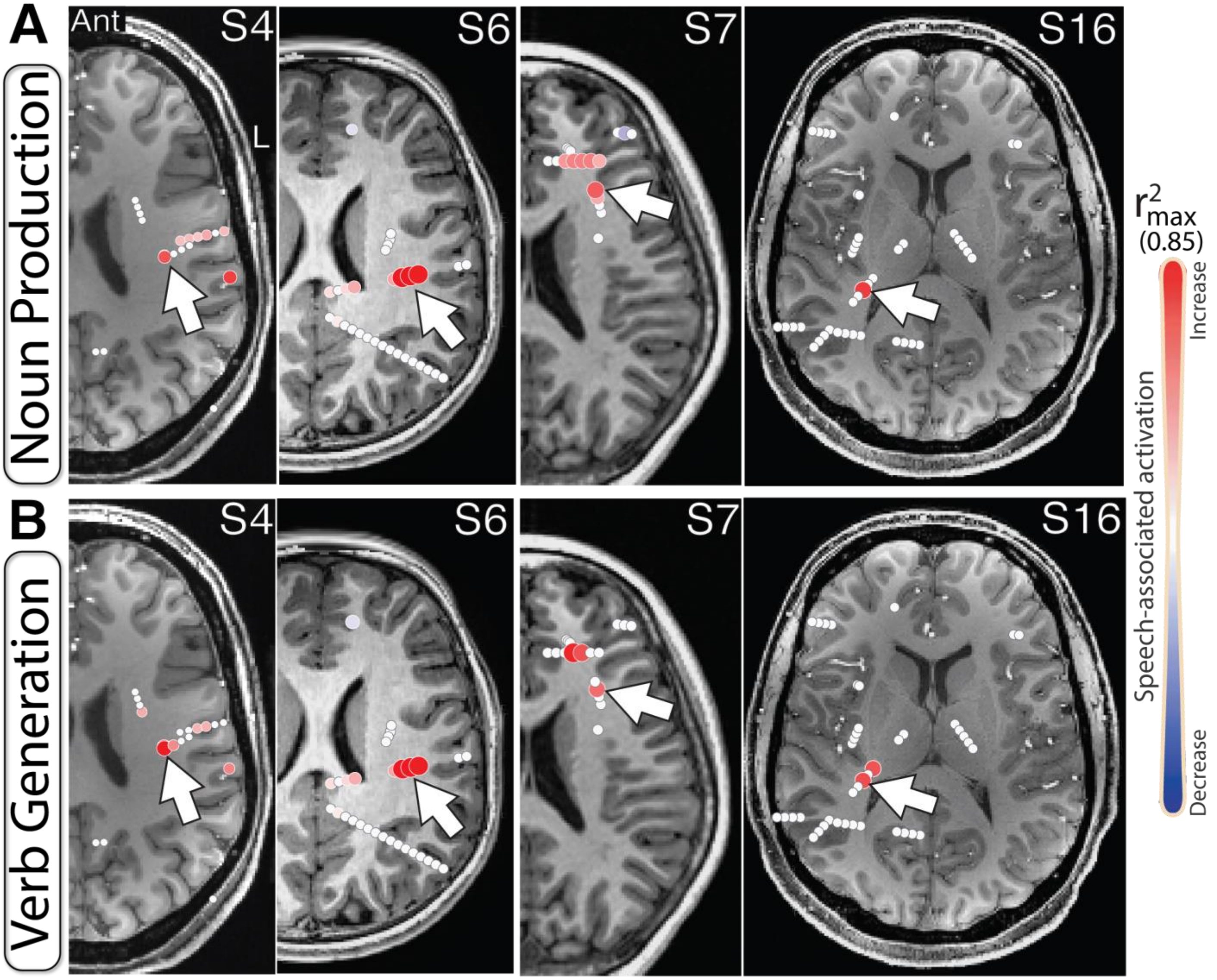


**Supplementary Figure 26. White matter activity during speech tasks. A.** T1 axial slices showing activity maps demonstrating broadband power shifts in white matter (white arrow) during a noun production task. **B.** As in A, but during a verb generation task.

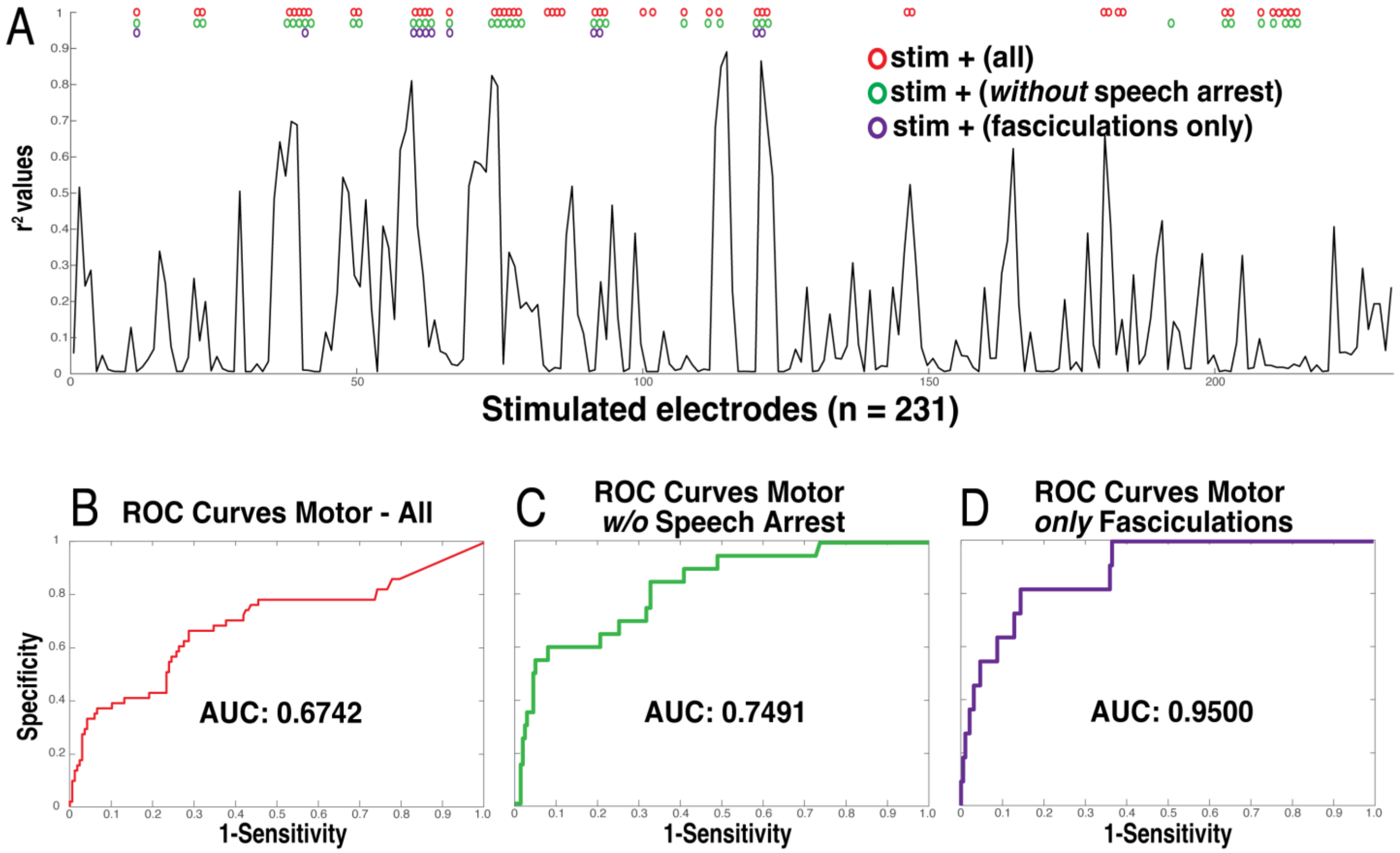


**Supplementary Figure 27. Summary Receiver Operating Characteristic (ROC) Curves Across Varying Thresholds of Positive Stimulation Results. A.** ROC curve assessing the performance of functional sEEG mapping relative to cortical stimulation (ground truth). **B.** Individual channel comparison of stimulation (circles) and functional (curve) sEEG mapping results. All stim responses that may be related to sensorimotor responses were considered positive. **C.** As in A, but when speech arrest responses were treated as negative responses. **D.** As in B, but when speech arrest responses were treated as negative responses. **E.** As in A, but when only stim sites with clear muscle fasciculation responses were treated as positive. **F.** As in B, but when only stim sites with clear muscle fasciculation responses were treated as positive.

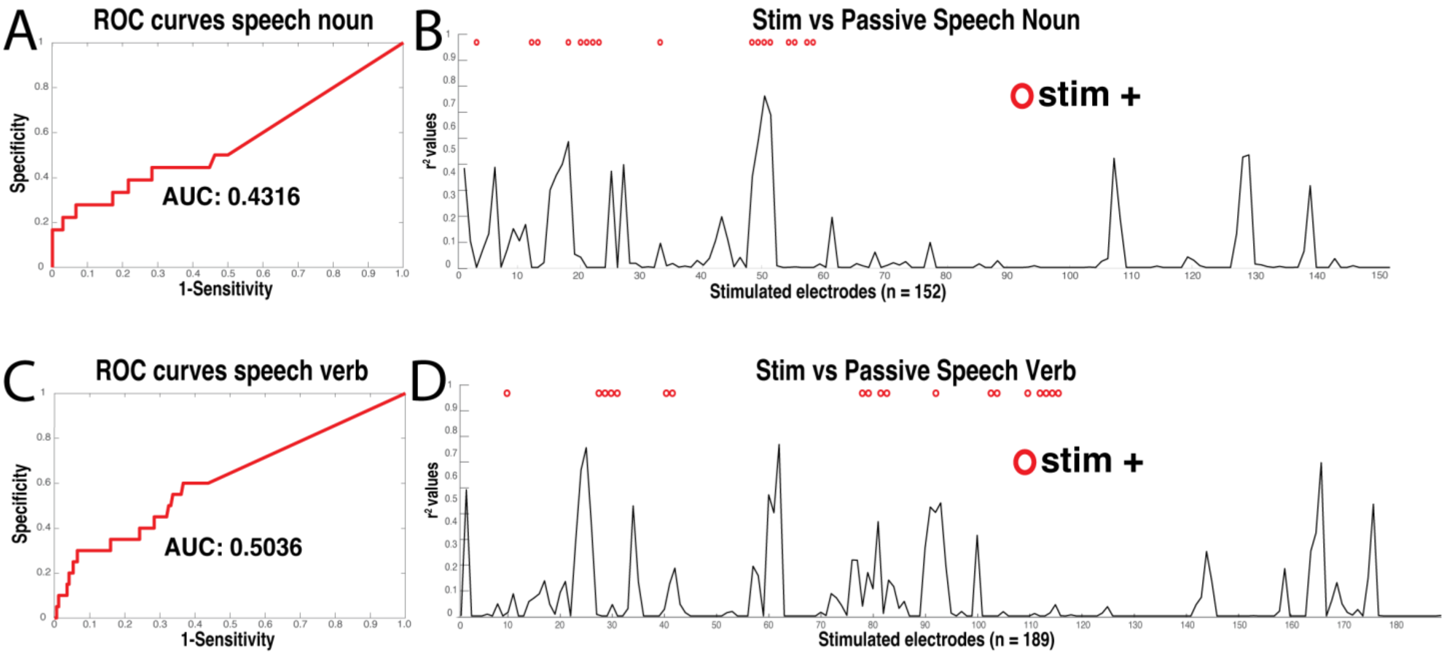


**Supplementary Figure 28. Summary Receiver Operating Characteristic (ROC) Curves Across Speech Tasks. A.** ROC curve assessing the performance of functional sEEG mapping relative to cortical stimulation (ground truth) during the noun production task. **B.** Individual channel comparison of stimulation (circles) and functional (curve) sEEG mapping results. All stim responses that may be related to speech responses were considered positive. **C.** As in A, but related to the verb generation task. **D.** As in B, but related to the verb generation task. Note that more patients performed the verb task than the noun allowing for more comparisons.

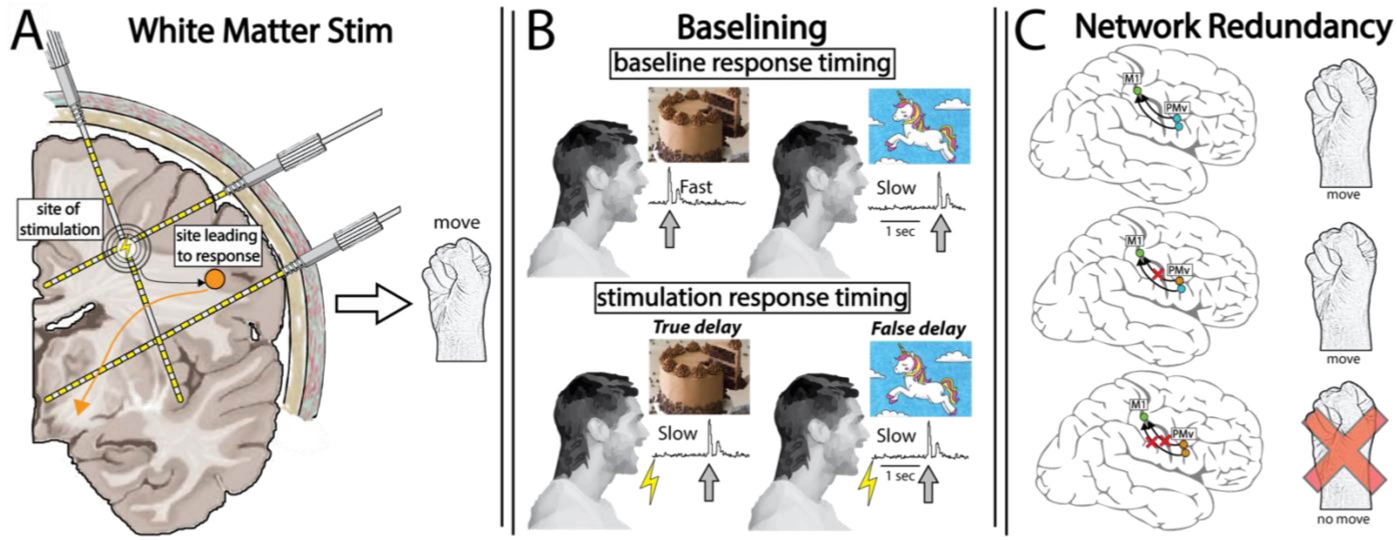


**Supplementary Figure 29. Potential Explanations for Differences Between Mapping Techniques. A.** White matter stimulation can lead to activation of distant cortex, leading to false positive stimulation results. This is most prevalent when bipolar stim pairs sit at a gray-white matter junction. **B.** Baseline data of the tasks carried out during stimulation mapping should be collected to identify true responses during stimulation mapping. Failure to do so may lead to false positive during stimulation mapping. **C.** Due to redundancy in speech and motor networks, stimulating a single node when other nodes can serve the same function will not lead to clinical responses. Failure to stimulate all redundant sites at once will lead to false negative stimulation results.
